# Obesity impairs therapeutic efficacy of mesenchymal stem cells by inhibiting cardiolipin-dependent mitophagy and intercellular mitochondrial transfer in mouse models of allergic airway inflammation

**DOI:** 10.1101/2021.11.27.470183

**Authors:** Shakti Sagar, Md. Imam Faizan, Nisha Chaudhary, Atish Gheware, Khushboo Sharma, Iqbal Azmi, Vijay Pal Singh, Gaurav Kharya, Ulaganathan Mabalirajan, Anurag Agrawal, Tanveer Ahmad, Soumya Sinha Roy

## Abstract

Mesenchymal stem cell (MSC) transplantation alleviates metabolic defects in diseased recipient cells by intercellular mitochondrial transport (IMT). However, the effect of host metabolic conditions on MSCs in general, and IMT in particular, has largely remained unexplored. This study has identified a molecular pathway that primarily governs the metabolic function and IMT of MSCs. We found underlying mitochondrial dysfunction, impaired mitophagy, and reduced IMT in MSCs derived from high-fat diet (HFD)-induced obese mice (MSC-Ob). Mechanistically, MSC-Ob failed to sequester their damaged mitochondria into LC3-dependent autophagosomes due to decrease in mitochondrial cardiolipin content, which we propose as a putative mitophagy receptor for LC3 in MSCs. Functionally, MSC-Ob exhibited diminished potential to rescue metabolic deficits and cell death in stress-induced epithelial cells. In a small molecule screen, we found pyrroloquinoline quinone (PQQ) as a regulator of mitophagy and IMT. Long-term culture of MSC-Ob with PQQ (MSC-Ob^PQQ^) restored cardiolipin content and sequestration of mitochondria to autophagosomes with concomitant activation of mitophagy. Upon co-culture, MSC-Ob^PQQ^ rescued cell death in stress-induced epithelial cells by enhancing IMT. The beneficial effect of PQQ was also evident in MSCs derived from human subjects in an in vitro model. In two independent mice models, the transplantation of MSC-Ob^PQQ^ restored IMT to airway epithelial cells, improved their mitochondrial metabolism and attenuated features of allergic airway inflammation (AAI). However, unmodulated MSC-Ob failed to do so. In summary, we uncover the molecular mechanism leading to the therapeutic decline of obese-derived MSCs and highlight the importance of pharmacological modulation of these cells for therapeutic intervention.

## Introduction

MSCs are widely being explored as promising cell-based therapies for several human diseases (*1, 2*). Clinical trials are underway to explore MSCs for the treatment of complex lung disorders (*2, 3*). A recent addition is the successful early phase clinical trial of MSCs to lower mortality due to COVID-19 (*4*). The beneficial effect of MSCs is attributed to their intrinsic immunomodulatory activity, generally mediated by their paracrine secretions (*5–7*). Over the last decade, a new paradigm of MSC therapeutics has emerged which is based on their unique ability to donate functional mitochondria to metabolically compromised recipient cells (*8–12*). The mitochondrial donation by MSCs to injured alveolar and bronchial epithelial cells alleviates acute lung injury and inflammatory airway diseases (*8, 13, 14*), independent of their immunomodulatory cytokine secretion and regenerative potential (*8*). On the flip side, mitochondrial donation by MSCs enhances bioenergetics in cancer cells (*15–17*) and increases their drug-resistance. This mitochondrial donation potential is also retained by MSCs derived from induced pluripotent cells (iPSCs), which alleviate asthmatic features and cigarette smoke-induced COPD (*18, 19*). Further, MSCs donate mitochondria directly to immune cells to modulate host cell immune response and alleviate tissue inflammation (*20*). Nevertheless, most of the studies exploring the therapeutic efficacy of MSCs have relied on the cells obtained from disease-free conditions. Induction of cell stress in MSC drastically reduces their intercellular mitochondrial donation capacity and compromises their therapeutic efficacy (*10*). Accumulating evidence now show that MSCs derived from disease-specific animal models and human patients exhibit impaired bioenergetics and altered mitochondrial quality control (MQC), which may compromise their long-term therapeutic efficacy (*21–25*).

Human adipose derived stem cells (ASCs) from aged individuals display oxidative stress, reduced multilineage proliferation rate, and decline in differential potential with a concomitant increase in senescence phenotype (*26, 27*). Similarly, ASCs derived from human orbital fat show senescent phenotype, reduced differential potential, and impaired stemness properties (*28*). These age-related changes markedly reduce the number of functional MSCs, which further limits their clinical application (*29*). Most notably, these changes are associated with a decline in mitochondrial function and elevated oxidative stress (*30*). Besides donor age, mitochondrial dysfunction is also observed in MSCs obtained from animal models of metabolic syndrome (MetS) and human patients with obesity, diabetes, MetS, and aging (*31, 32*). Obese condition also has a pronounced effect on MSC immunomodulatory activity, stem cell property, and therapeutic efficacy (*23, 33*). This has important clinical implications for autologous MSC treatment in such patients, especially for indications related to complications of metabolic disorders. Here we examine whether intercellular mitochondrial donation capacity and therapeutic efficacy of MSC from obese subjects is compromised. Further, we examine whether mitochondria-targeted therapy can reverse such defects, enabling autologous treatment.

Generally, dysfunctional mitochondria are selectively removed from the cells by mitophagy that reduces the accumulation of damaged mitochondria to prevent cell death (*34, 35*). Mitophagy is normally followed by mitochondrial biogenesis to ensure an adequate number of functional mitochondria through the mitochondrial quality control (MQC) pathway (*36*). Under certain disease conditions, mitochondrial dysfunction may not always be accompanied by mitophagy, which results in metabolic decline and eventually cell death (*37*). Classical mitophagy involves stabilization of mitochondrial serine/threonine-protein kinase PINK1 on the outer membrane of depolarized mitochondria and recruitment of cytosolic E3-ubiquitin-protein ligase Parkin to these depolarized mitochondria. Though, alternate Parkin independent mitophagy mediated by BNIP3L/NIX and FUNDC1 exists in certain cells, (*38–40*) recent studies suggest that MSCs mostly utilize the classical PINK1/Parkin pathway to clear the damaged mitochondria (*41, 42*). Conditions such as senescence, obesity, and type II diabetes mitigate autophagy/mitophagy in MSCs, resulting in the accumulation of dysfunctional mitochondria and compromised beneficial effect (*43–46*). However, the mechanism of mitochondrial dysfunction and the role of mitophagy in determining the therapeutic potential of MSCs has largely remained unexplored.

Here, we have systematically evaluated the role of mitophagy in MSCs derived from the (HFD)-induced obese mice and human MSCs and evaluated their therapeutic potential in vitro and in two independent mouse models of allergic asthma. Our findings uncover previously unknown mechanism that determine the mitophagy, IMT and thereupon the therapeutic efficacy of MSCs. Importantly, we have shown that the impaired therapeutic potential of MSC-Ob is reversible with a small molecule PQQ, which restores mitophagy and IMT by enhancing sequestration of dysfunctional mitochondria to autophagosomes.

## Results

### MSCs derived from obese mice show reduced intercellular mitochondrial transport and accumulation of dysfunctional mitochondria

We developed an 18-week-old (HFD)-induced obese mice model as described by us previously (Fig. 1A) (*47*). An increase in body weight (44.2±0.68 in obese vs. 24.9±00.48 in lean) and changes in biochemical profile confirmed the obesity features in these mice (Fig. S1). To determine whether MSCs derived from HFD mice maintain stem cell property, we harvested cells from lean (MSC-L) and HFD mice (MSC-Ob) and analyzed them for expression of key stem cell markers (positive: Sca1, CD44; negative: CD11b). Both the groups had similar expression of the stem cell markers (Fig. S2A). However, MSC-Ob showed a trend towards increase in cell death and cellular senescence but decrease in the rate of cell proliferation (Fig. S2B-E). Thus, MSCs derived from (HFD)-induced obese mice, despite retaining the stem cell markers, show alteration in overall cell survival in the culture.

**Fig. 1.**
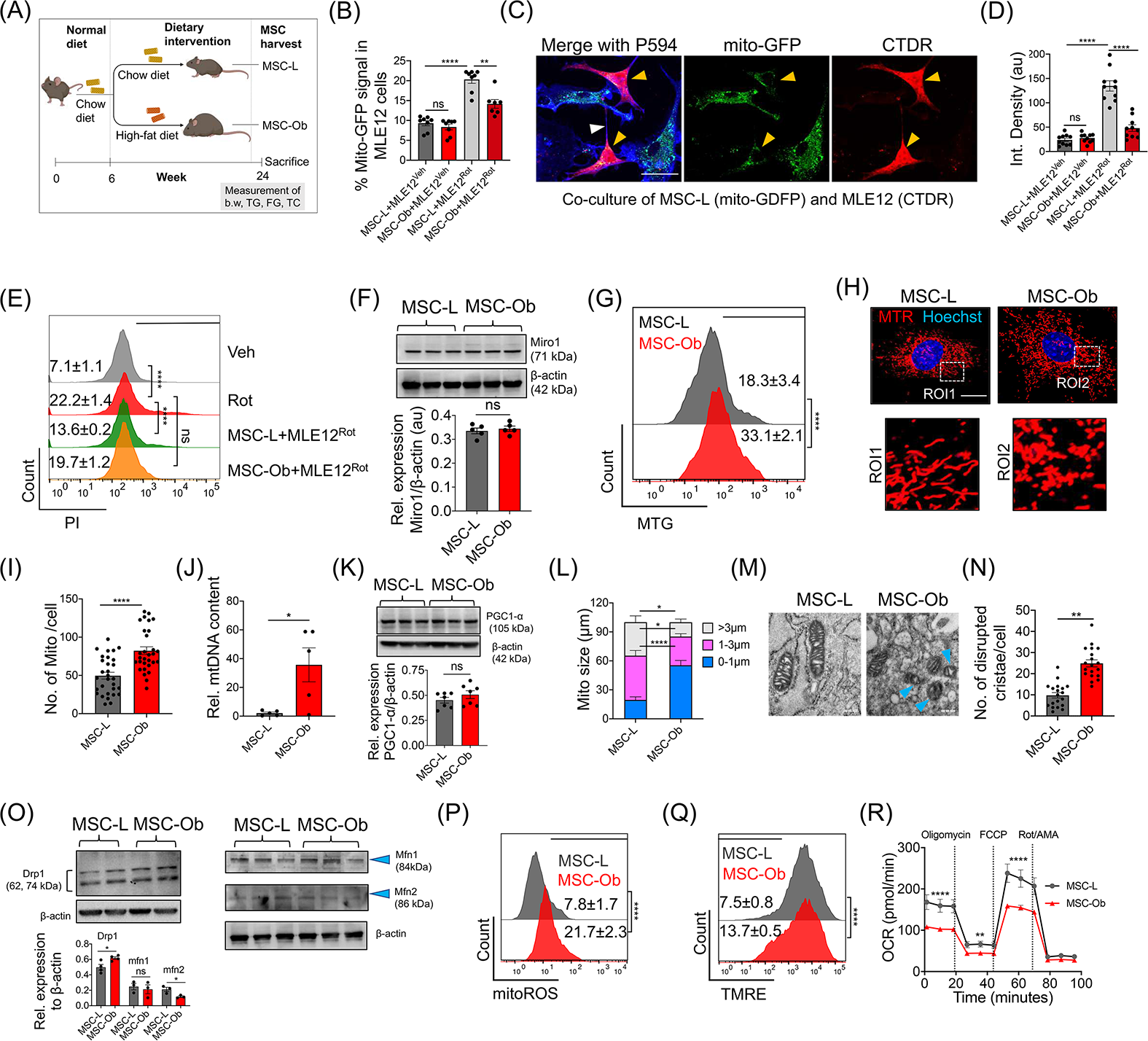
Obese-derived MSCs exhibit reduced intercellular mitochondrial transport due to mitochondrial dysfunction: **(A)** Schema of the development of diet-induced obesity model. The MSCs were harvested from mice fed a high-fat diet for 18 weeks (MSC-Ob) and control mice, which received a regular diet (MSC-L). The mice were subjected to measurements of body weight (b.w.), fasting glucose (FG), triglycerides (TG), and total cholesterol (TC) before sacrifice at week 24. (**B**) Flow cytometry analysis showing mitochondrial donation by GFP transduced MSCs (mito-GFP) to the vehicle (Veh), or rotenone (Rot) treated MLE-12 cells after 24 hrs of co-culture. (**C**) Representative images of mito-GFP-transduced MSC-L show mitochondria donation to Rot treated MLE12, which were stained with cell-tracker deep red (CTDR). Yellow arrowheads show the mito-GFP signal in MLE12 cells, and TNTs (white arrowhead) between the two cells visualized after fixing the cells and staining with phalloidin 594 (P594; blue). (**D**) The integrated density of mito-GFP signal quantified in CTDR positive MLE12 treated with Veh or Rot. (**E**) Representation of flow cytometry histograms showing mtROS levels in cell tracker green (CTG) stained MLE12 cells after co-culture with unstained MSCs. (**F**) Immunoblot of Miro1 expression in total cell lysate and the corresponding densitometric analysis *(below panel)*. (**G**) The mitochondrial mass measured by flow cytometry in MSC-L and MSC-Ob after staining with mitotracker green (MTG). (**H**) Representative images of MSC-L and MSC-Ob after staining with mitotracker red (MTR) and Hoechst (blue). (**I**) mitochondrial cell number was calculated by staining the cells with MTR and imaging analysis and presented as number of mitochondria per cell. (**J**) RT-qPCR analysis showing mtDNA content in MSC-L and MSC-Ob. (**K**) Immunoblot of PGC-1α and β-actin in cell lysate from MSC-L and MSC-Ob along with densitometric analysis (*below panel*). (**L**) Mitochondria size was calculated in images obtained after staining the cells with MTR. (**M, N**) EM images show cristae disruption, more pronounced in MSC-Ob (blue arrowheads) and quantitive analysis per 100 cells. (**O**) Immunoblot and densitometric analysis of Drp1, mfn1, mfn2, and β-actin. (**P, Q**) graphical representation of flow cytometry data showing mtROS in cells stained with mitoSOX and percentage depolarized mitochondria in cells stained with TMRE. (**R**) Measurement of oxygen consumption rate (OCR) in MSC-L and MSC-Ob under basal and various mitochondrial complex inhibitor treatments. Data is shown as Mean±SEM with n ≥ 3. \*\*\*\**P* < 0.001; \*\*\**P* < 0.005; \*\**P* < 0.01; \**P* < 0.05; ns (non-significant). Scale bars: C: 20 µm; D: 10 µm; M: 0.1 µm.

It has largely remained elusive as to what extent MSCs derived from diseased, particularly from obese patients retain therapeutic efficacy. To probe the same, we used a co-culture system to model mitochondrial donation, using mito-GFP expressing MSCs as donors and stressed mouse lung epithelial cells (MLE12) as mitochondria recipients. MLE12 cells were first treated with either vehicle (Veh) or rotenone (Rot) (to induce mitochondrial dysfunction) for 12 hours (hrs) and then stained with cell tracker deep-red before co-culture. After 24 hrs of co-culture, we performed flow cytometry analysis to measure the percentage of MLE12 cells expressing mito-GFP – an indicator of mitochondrial uptake. A substantial decrease in intercellular mitochondrial transport from MSC-Ob to the MLE12^Rot^ cells was observed, compared to that from MSC-L (Fig. 1B). We found a similar trend using imaging studies followed by quantitative analysis (Fig. 1C-D). This decline in the mitochondrial donation by MSC-Ob was associated with their diminished potential to rescue cell death in MLE12^Rot^ cells (Fig. 1E).

We have previously reported that Miro1 regulates the intercellular transfer of mitochondria by MSCs (*8*). To examine whether the decrease in mitochondrial donation was due to the differential expression of Miro1, we measured its expression at mRNA and protein levels. However, MSC-L and MSC-Ob did not significantly differ in the Miro1 expression (Fig. 1F and S 3A), which is consistent with our previous study that endogenous Miro1 expression does change during epithelial cell stress (*8*). Similarly, we did not find any significant changes in tunneling nanotube (TNT) formation, which are the primary mediators of IMT (Fig. S3B, C) (*48*).

The decrease in IMT can also attribute to enhanced mitochondrial turn-over or reduced mitochondrial biogenesis. On the contrary, we found a distinctly increased mitochondrial mass, mtDNA content, and mitochondrial number per cell in MSC-Ob, as measured by flow cytometry, imaging analysis, and RT-qPCR (Fig. 1G-J). The increase in mitochondrial mass was not due to mitochondrial biogenesis as reflected by unaltered PGC-1α expression (Fig. 1K). Notably, MSC-Ob displayed smaller and punctate mitochondrial structure with disrupted cristae than the regular tubular network seen in MSC-L (Fig. 1L-N). In line with changes in mitochondrial shape, we found trend towards increase in Drp1 (mito-fission protein) with a concomitant decrease in Mfn2 (mito-fusion protein) expression (**Fig. O**). These mitochondrial shape and form changes corroborate increased mitochondrial ROS (mtROS) levels, reduced mitochondrial membrane potential (ΔΨm), and decreased bioenergetic flux (Fig. 1P-R). Prominently, ATP levels, basal respiration, maximal respiration, and spare respiratory capacity were significantly reduced (Fig. S4). Altogether, these results indicate that MSCs derived from obese mice exhibit reduced intercellular mitochondrial donation, apparently due to the accumulation of dysfunctional mitochondria.

### MSCs derived from obese mice show altered mitochondrial sequestration to autophagosomes

Accumulation of dysfunctional mitochondria is generally attributed to their reduced clearance from the cells by mitophagy (*49*). Therefore, we looked at the conventional pathway of mitophagy in MSCs, which involves PINK1 and Parkin. MSCs were treated with FCCP (5 and 10 µM) to induce mitochondrial depolarization, and mitochondrial fractions were prepared and subjected to immunoblotting for endogenous PINK1 and Parkin. The quality of mitochondrial fraction was confirmed by checking for the expression of mitochondrial marker and the absence of cytosolic marker (Fig. S5A). As shown in Fig. 2A, FCCP treatment induced accumulation/stabilization of PINK1 in the mitochondrial fractions derived from MSC-L. Surprisingly, MSC-Ob showed inherent stabilization of PINK1, with no further increase upon depolarization (Fig. 2A). Parkin showed a similar pattern of expression (Fig. 2B) and these results were confirmed by immunofluorescence followed by colocalization analysis (as calculated by Mander’s coefficient) (Fig. 2C, D). These results are thus consistent with the notion that depolarized mitochondria stabilize PINK1 and recruit cytosolic Parkin (*34, 50*). This pattern of expression seen in MSC-Ob indicates their inherently depolarized mitochondria, as shown in *Fig. 1Q*.

**Fig. 2.**
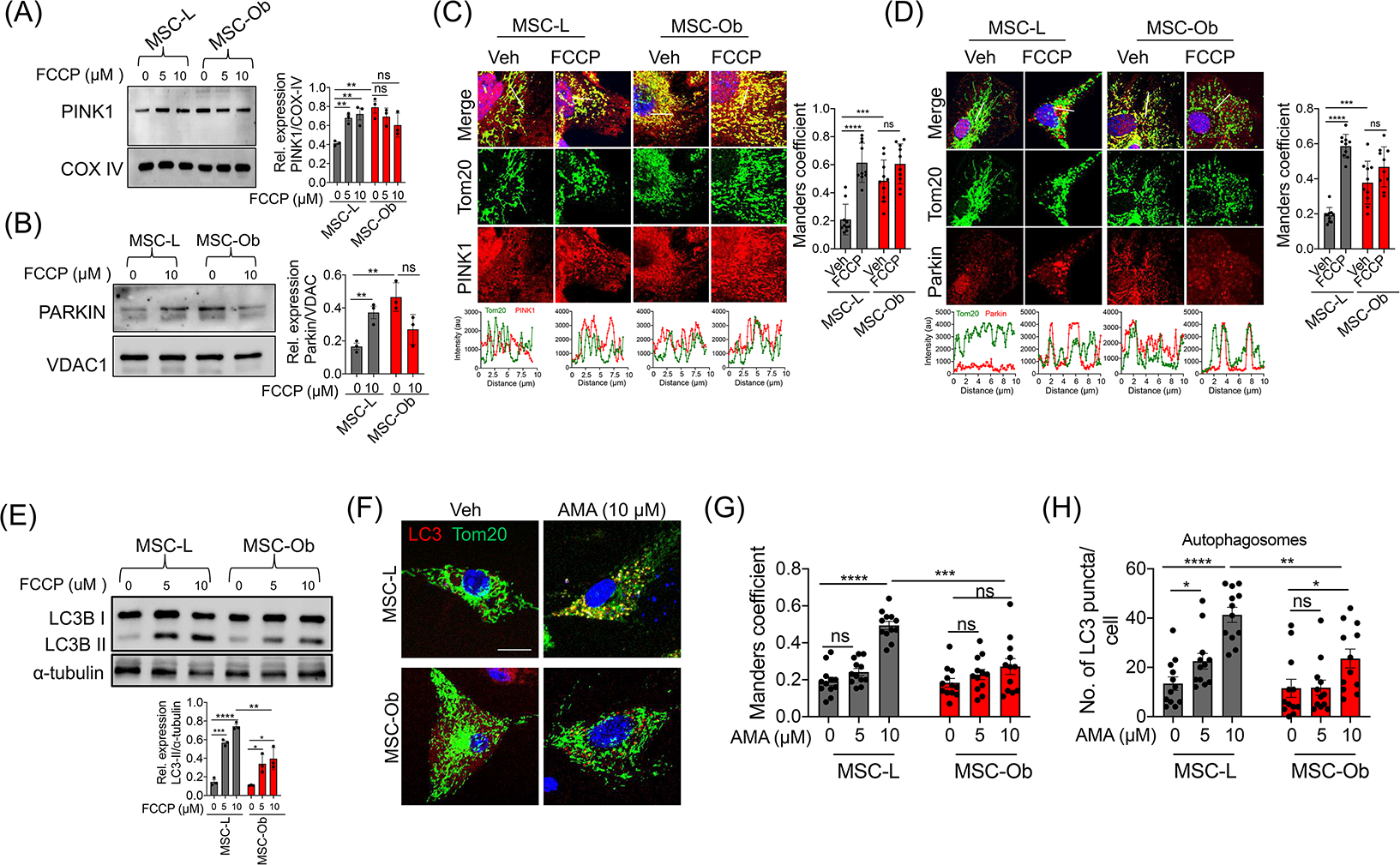
MSC-Ob exhibits impaired mitophagy and reduced activation of LC3-dependent autophagosomes: (**A**) Immunoblot of PINK1 in the mitochondrial extracts prepared under DMSO (0) and FCCP (5µM and 10µM) treatment for 1 hr with densitometric analysis (*right panel*). (**B**) Similarly, immunoblot showing the expression of Parkin. (**C**) Representative images of cells stained for endogenous PINK1 (red) and Tom20 (green) treated with Veh or FCCP. Line scans show colocalization (below panel) between PINK1 and Tom20. Images from *(C)* were analyzed to determine Mander’s coefficient (right panel), which indicates the extent of colocalization between mitochondria (green) and PINK1 (red). (**D**) Similarly, images of Parkin’s immunofluorescence and line scan analysis *(below panel)* and Mander’s coefficient. (**E**) Immunoblot of LC3 in MSCs treated with DMSO (0) or FCCP (5µM and 10µM) for 1 hr. The lower panel shows the densitometry analysis. (**F**) Representative images of LC3 (red) and Tom20 (green) treated with DMSO (Veh) or 10µM antimycin (AMA) for 1 hr. (**G**) Mander’s coefficient representing the colocalization between LC3 and Tom20. (**H**) Quantitation of LC3 puncta per cell, which represents the autophagosome formation in images from panel *G*. \*\*\*\**P* < 0.001; \*\*\**P* < 0.005; \*\**P* < 0.01; \**P* < 0.05; ns (non-significant). Scale bars: A: 50 µm; B: 10 µm; D, E: 20 µm.

We next assessed the role of essential proteins implicated in autophagosome formation. Depolarized mitochondria are generally sequestered to the autophagosomes and thereupon to lysosomes for clearance (*34, 50*). The autophagosome formation begins with the maturation of diffused LC3 to the puncta forming LC3-II, which is also considered as a marker for autophagosomes. First, we performed immunoblotting to probe any changes in the maturation of LC3-II. As shown in Fig. 2E, FCCP treatment induced LC3-II formation in MSC-L, whereas MSC-Ob had significantly lower levels. Next, we evaluated LC3-dependent autophagosome formation and LC3 colocalization with mitochondria under un-induced and chemically-induced depolarization conditions. We used a mild depolarizing agent antimycin A (AMA), which induces mitochondrial dysfunction by enhancing mtROS production (*51*). As expected, we found LC3 puncta formation in MSC-L and their colocalization with mitochondria upon depolarization. On the contrary, MSC-Ob showed significantly reduced colocalization with a concomitant decrease in the autophagosome number (Fig. 2F-H **and** S5B, C). We also assessed the effect of LC3 overexpression on autophagosome formation and mitochondrial sequestration. However, we did not find any significant changes in LC3 maturation and colocalization with mitochondria in MSC-Ob after LC3 overexpression (Fig. S5D). These results thus indicate that reduced sequestration of dysfunctional mitochondria into the autophagosomes contributes to their accumulation in MSC-Ob.

### Increased lysosomal content in MSC-Ob does not correlate with improved mitophagy

To examine the involvement of the general autophagy pathway, we assessed the expression of crucial autophagy regulatory proteins. However, we did not find any significant changes in the expression of Atg5, Atg7, total and phosphorylated Beclin1 (Fig. S6A-C). Similarly, using an autophagy pathway specific RT-qPCR array, we did not find differential expression in their transcript levels (Fig. S6D). These results suggest that the expression of general autophagy proteins remains unaltered in MSC-Ob, despite significant alterations in the mitophagy pathway.

The autophagosomes eventually fuse with the lysosomes to form the autophagolysosome. Previous studies have reported impaired mitophagy during lysosomal abnormalities (*52*). We thus assessed the status of lysosomes by evaluating the expression of late endosome/lysosome marker LAMP1 and lysosomal content. Surprisingly, we observed a significant increase in basal LAMP1 expression in MSC-Ob. On the contrary, MSC-L showed increased expression only upon depolarization (Fig. 3A, B). Additionally, we performed live-cell imaging of cells stained with lysotracker deep red (LTDR) followed by image analysis. As shown in Fig. 3C, an increase in the LTDR staining in MSC-Ob was observed. These results thus suggest that MSC-Ob has inherently higher lysosomal content.

**Fig. 3.**
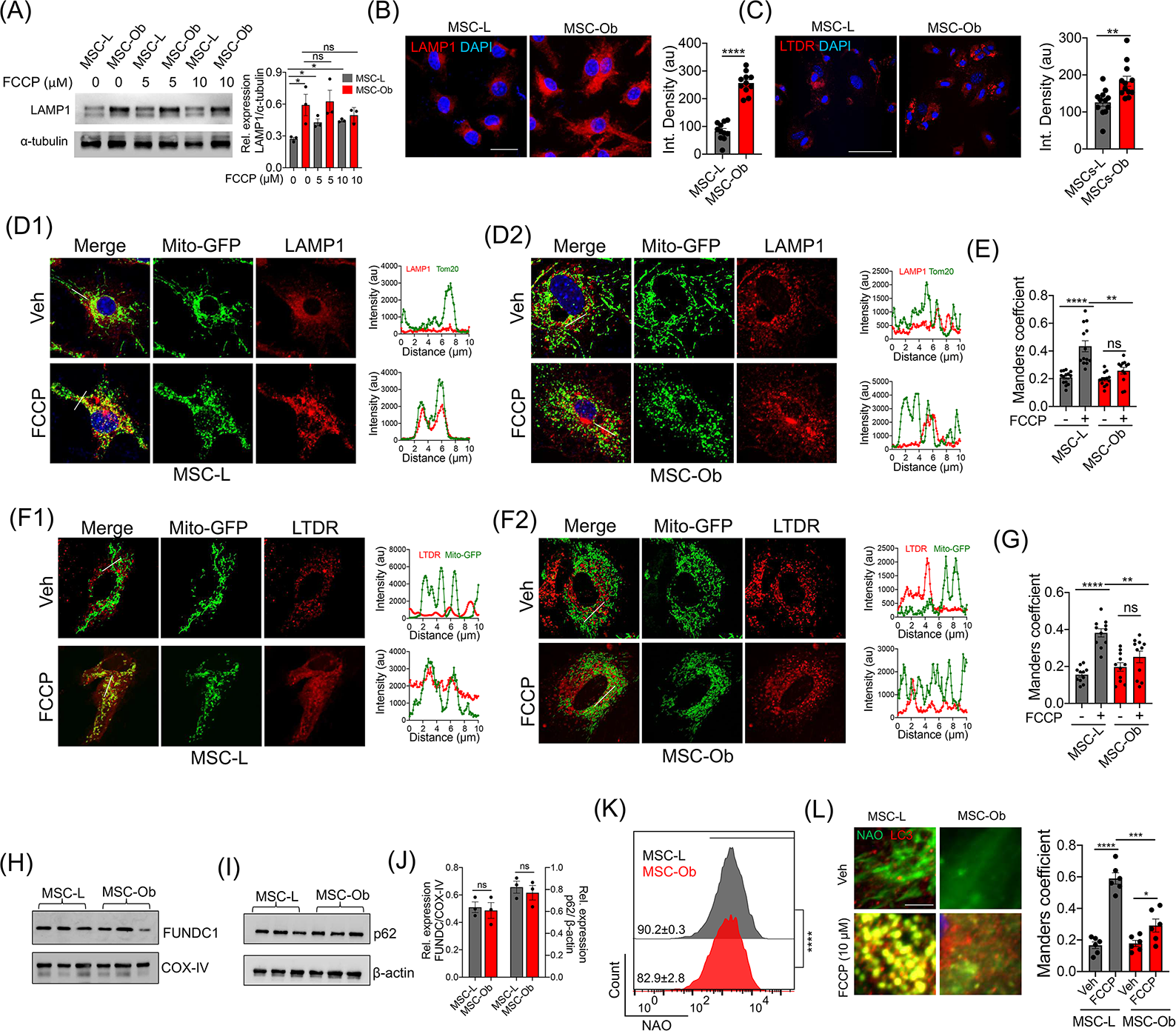
Mitochondrial sequestration into autophagosomes is inhibited by reduced cardiolipin content rather changes in lysosomal function: **(A)** LAMP1 expression in MSCs treated with DMSO or FCCP for 2 hrs before the total protein lysates were prepared for immunoblotting. The right panel shows the densitometry analysis of the blots. (**B**) Representative images of LAMP1 in MSCs stained with anti-LAMP1 antibody (red) and DAPI (blue). The right panel shows the image analysis data plotted as integrated density. (**C**) MSCs were live stained with lysotracker deep-red (LTDR) and imaged in the presence of Hoechst stain (blue). The images *(C)* were quantified and represented as integrated density. (**D**) Representative images of MSC-L (D1) and MSC-Ob (D2) transduced with mitochondrial-targeted GFP (mito-GFP) and treated with DMSO or FCCP for 2 hrs. After fixation, the cells were stained with LAMP1 (red) and DAPI. *Right panels* show the line scans of the images indicating the extent of colocalization. (**E**) Mander’s coefficient showing the degree of colocalisation between LAMP1 and mitochondria. (**F**) Similarly, the images of MSC-L *(F1)* and MSC-Ob *(F2)* along with the line scans. (**G**) Image analysis *(D)* to determine the Mander’s coefficient between lysosomes and mitochondria. MSCs after transduction with mito-GFP (green) were treated with DMSO or FCCP and further stained with lysotracker deep-red (LTDR; red). Line scans indicate the extent of overlap between LAMP1 and mitochondria in the region selected *(F).* (H, I) Expression of FUNDC1, COX-IV, p62 and β-actin as revealed by immunoblotting of MSCs treated with DMSO (0) or FCCP (5µM and 10µM). (**J**) The corresponding densitometry analysis. (**K**) Histograms showing the cardiolipin content as measured by staining the cells with NAO. (**L**) Representative images showing the colocalisation of LC3 (red) with NAO (green). Cells were treated with Veh or FCCP (10 µM) for 2 hrs. (*Right panel*) Corresponding Mander’s coefficient of the images *(L)* showing the colocalisation between NAO and LC3. Mean±SEM. Mean±SEM. \*\*\*\**P* < 0.001; \*\*\**P* < 0.005; \*\**P* < 0.01; \**P* < 0.05; ns (non-significant). Scale bars: B: 50 µm; C: 100 µm; D, F: 10 µm; L: 5 µm.

Given the fact that MSC-Ob have increased lysosomal content and an alternative autophagy pathway exists wherein depolarized mitochondria are directly taken up by the endosomal pathway for clearance (*53, 54*), we investigated whether increase in lysosomal content primes these cells to use this alternative for mitochondrial clearance. To explore this possibility, we transduced the cells with lentiviral mito-GFP and subsequently stained them with LAMP1 and LTDR, respectively. MSC-Ob did not show any significant changes in the association of mitochondria with LAMP1 (Fig. 3D) or LTDR (Fig. 3F) with or without FCCP treatment, as opposed to MSC-L. Mander’s coefficient also revealed reduced colocalization of mitochondria with LAMP1 (0.43±0.03 MSC-L FCCP vs. 0.24±0.03 MSC-Ob FCCP) with a similar trend observed in LTDR (0.38±0.03 MSC-L FCCP vs. 0.25±0.02 MSC-Ob FCCP) (Fig. 3E, G). To check whether MSC-Ob has a general defect in the autophagic flux, we used a control autophagy assay based on p62/SQSTM1 turnover (*55*). Autophagy was induced in the cells by starvation. Consistent with the results above (Fig. 2), autophagosome formation was induced in MSC-Ob only upon starvation, albeit lower than MSC-L. Notably, MSC-Ob showed relatively lower p62 colocalization with LC3 and LAMP1 than MSC-L (Fig. S7). Together, these results illustrate that an increase in lysosomal content observed in MSC-Ob does not correlate with clearance of dysfunctional mitochondria.

### MSC-Ob and hMSC^FFA^ display reduced cardiolipin content which is a putative mitophagy receptor for LC3

For activation of downstream mitophagy, LC3-containing autophagosome binds to the depolarized mitochondria via mitophagy receptors. Altered expression of these receptors impairs sequestration of damaged mitochondria into autophagosomes (*36, 56*). We thus evaluated the expression pattern of these receptors. While FUNDC1 and p62 expression did not significantly differ between MSC-L and MSC-Ob, a significant decrease in cardiolipin (stained with NAO) content was observed (Fig. 3H-K). We also observed that MSC-Ob cardiolipin-stained mitochondria showed reduced colocalization with LC3 upon depolarization (Fig. 3L **and** S8). Our findings (including *Fig. 1M*) are consistent with previous reports that decrease in cardiolipin causes cristae disruption and impairs mitochondrial clearance due to reduced interaction with LC3-containing autophagosomes (*57, 58*). Thus, these results demonstrate that obese-derived MSCs with underlying mitochondrial dysfunction and reduced cardiolipin content fail to sequester their depolarized mitochondria into the autophagosomes.

To correlate our findings in human MSCs, we also evaluated the impact of free fatty acids (FFA) on mitochondrial health, cardiolipin content and mitophagy, in an in vitro model of a human (h) MSC^FFA^. Cells were harvested from normal healthy subjects with no known history of MetS or obesity and characterized for stem cell markers (Fig. S9A). The in vitro model depicting features of MetS or obesity was developed by treating the cells with FFA for 24 hrs as described earlier (*59*). Upon culturing in FFA, hMSCs showed increased mtROS, pronounced punctate mitochondrial morphology, and increased mitochondrial mass (Fig. S9B-D). Additionally, hMSC^FFA^ showed a significant reduction in cardiolipin content (Fig. S9E). Upon depolarization, LC3 colocalised with cardiolipins to a much higher extent in Veh than hMSC^FFA^ (Fig. S9F). Collectively, these findings reveal that like MSC-Ob, hMSC^FFA^ also exhibit accumulation of dysfunctional mitochondria and reduced cardiolipin content which may impede the clearance of defective mitochondria.

### PQQ treatment restores impaired mitophagy by increasing sequestration of damaged mitochondria to autophagosomes

From the results above, it is evident that MSC-Ob has dysfunctional mitochondria with associated impaired mitophagy. We thus hypothesized that restoring mitochondrial health or autophagy will increase clearance of depolarized mitochondria. We choose drugs that directly or indirectly regulate mitochondrial health, mitophagy and autophagy. As an assay outcome, we used endogenous Tom20 and LC3 as markers to find the effect of these small molecules in inducing sequestration of mitochondria into autophagosomes. Images were taken and subjected to image analysis to determine the extent of colocalization between LC3 and mitochondria. Interestingly, we found relatively higher colocalization by pyrroloquinoline quinone (PQQ) than other molecules tested (Fig. 4A). Earlier studies have reported that PQQ attenuates mtROS and improves mitochondrial health, and increases autophagy (*60–62*), so based on these rational we choose PQQ in subsequent experiments to explore its role in alleviating mitochondrial health of MSC-Ob. Initially, we tested various time points of PQQ to find the optimal dose by looking at its effect in lowering the mtROS. We found that chronic treatment (6 doses at an interval of 48 hrs) of 30 µM PQQ (hereafter referred to as PQQ) significantly reduced mtROS in MSC-Ob^PQQ^ as compared to any other dose and time point tested (Fig. 4B **and** S10B). PQQ treatment also restored mitochondrial, ΔΨm, and enhanced mitochondrial bioenergetics (Fig. 4C-E). Moreover, MSC-Ob^PQQ^ showed more elongated and tubular-shaped mitochondria than punctate form, with a concomitant decrease in mitochondrial mass and restoration of damaged cristae, suggesting clearance of the damaged mitochondria (Fig. 4F-H **and** S10C, D).

**Fig. 4.**
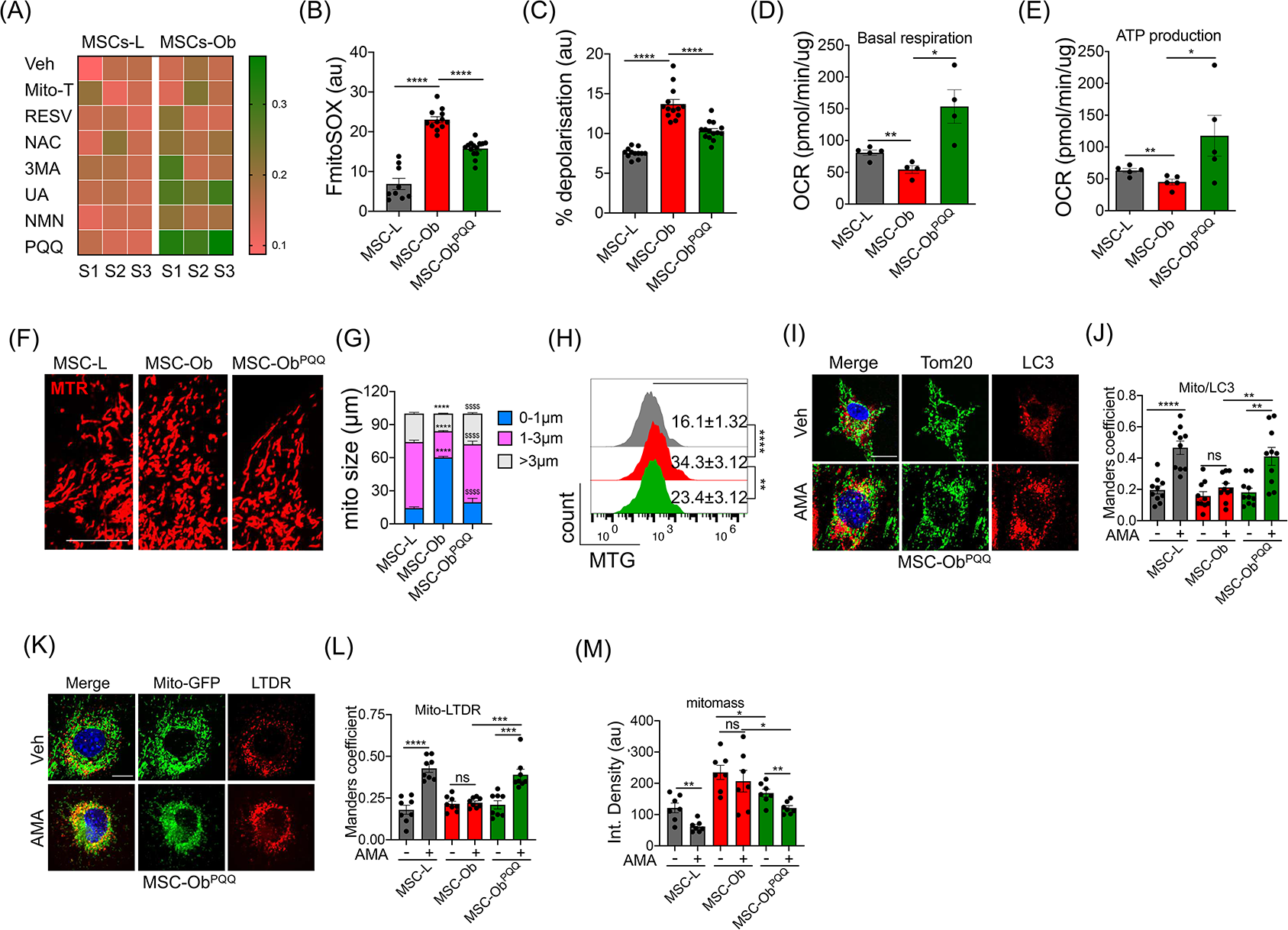
MSC-Ob cultured in PQQ restores structure and function of mitochondria and their sequestration to autophagosomes: (**A**) MSCs were treated with DMSO (Veh), Mito-TEMPO (Mito-T), Resveratrol (RESV), N-acetyl cysteine (NAC), 3-methyladenine (3MA), Urolithin A (UA), nicotinamide mononucleotide (NMN), and Pyrroloquinoline quinone (PQQ) for 48 hrs before fixing the cells. The cells were stained for endogenous LC3 (red) and Tom20 (green) and analyzed to determine the Mander’s coefficient represented as heat-map. (**B**) Bar graph representation of flow cytometry analysis of mtROS in MSCs treated with PQQ (chronic 30 µM treatment of 6 doses for 10 days). The cells were stained with mitoSOX red. (**C**) Similarly, cells were stained with TMRE to determine the degree of mitochondrial depolarization. (**D, E**) OCR reflects ATP production and basal respiration. (**F**) Representative images of MSCs stained with mitotracker red (MTR) showing mitochondrial morphology. (**G**) Image quantitation *(F)* showing changes in mitochondrial size between different groups. (**H**) Representative flow cytometry histograms of mitotracker green (MTG) stained MSCs. (**I**) MSCs were treated with AMA (10µM) for 1 hr and then stained for endogenous LC3 (red) and Tom20 (green). (**J**) Mander’s coefficient was calculated in images from *(J)* to find the colocalization between Tom20 and LC3. (**K, L**) Similarly, cells were stained with lysotracker deep red (LTDR) and Tom20. (**M**) MSCs were treated with AMA (10µM) for 2 hr and stained with mitotracker green to find the mitochondrial turn-over, represented as integrated density. Mean±SEM. \*\*\*\**P* < 0.001; \*\*\**P* < 0.005; \*\**P* < 0.01; \**P* < 0.05; ns (non-significant). Scale bars: 10 µm.

We next assessed the effect of PQQ in the clearance of damaged mitochondria. A significant increase in the colocalization of mitochondria with LC3-autophagosomes was observed in MSC-Ob treated with PQQ (Fig. 4I-K). This effect of PQQ was corroborated with increase in the cardiolipin content and colocalisation with LC3 (Fig. S11A-C) with a similar trend observed in hMSC^FFA^ treated with PQQ (Fig. S11D-E). Similarly, we found that PQQ enhanced the recruitment of mitochondria to the lysosomes upon AMA treatment (Fig. 4L, M). Functionally, the effect of PQQ was evident from the clearance of chemically-induced depolarized mitochondria from MSC-Ob^PQQ^ but not from untreated MSC-Ob, which accumulate the damaged mitochondria (**Fig. N**). These results indicate that PQQ induces cardiolipin levels which enhances sequestration of dysfunctional mitochondria into LC3-containing autophagosomes.

### MSC-Ob^PQQ^ rescues mitochondrial damage in stressed epithelial cells by enhancing the intercellular transport of functional mitochondria

We and others have previously reported that donation of healthy functional mitochondria is integral to the therapeutic potential of MSCs, while a donation of dysfunctional mitochondria deteriorates the health of recipient cells (*8, 63*). Here, we asked whether MSC-Ob^PQQ^ can rescue epithelial cell apoptosis by donating functional mitochondria. As shown in Fig. 5A, in comparison to the untreated cells, PQQ treatment significantly increased IMT potential of MSC-Ob to damaged epithelial cells. This mitochondrial donation was associated with attenuation of mtROS and restoration of ΔΨm in recipient MLE12 cells (Fig. 5B, C). Further, the mitochondrial donation by MSC-Ob^PQQ^ restored mitochondrial shape, mitochondrial mass, and attenuated cell death in MLE12 cells (Fig. 5D-F). The effect of PQQ to restore IMT was found independent of its effect on Miro1 expression and TNT formation (Fig. 5G, H). These results thus suggest that mitochondrial health regulates IMT in MSCs. While partially depolarized mitochondria may still undergo IMT (*42*), severe depolarization significantly compromises this property. These findings are consistent with our previous findings that Rot-induced MSCs have impaired mitochondrial donation potential (*8*). Notably, the results presented here demonstrate that obesity-associated changes in MSC-Ob are reversible. Their therapeutic efficacy can be restored by enhancing sequestration of dysfunctional mitochondria into autophagosomes.

**Fig. 5.**
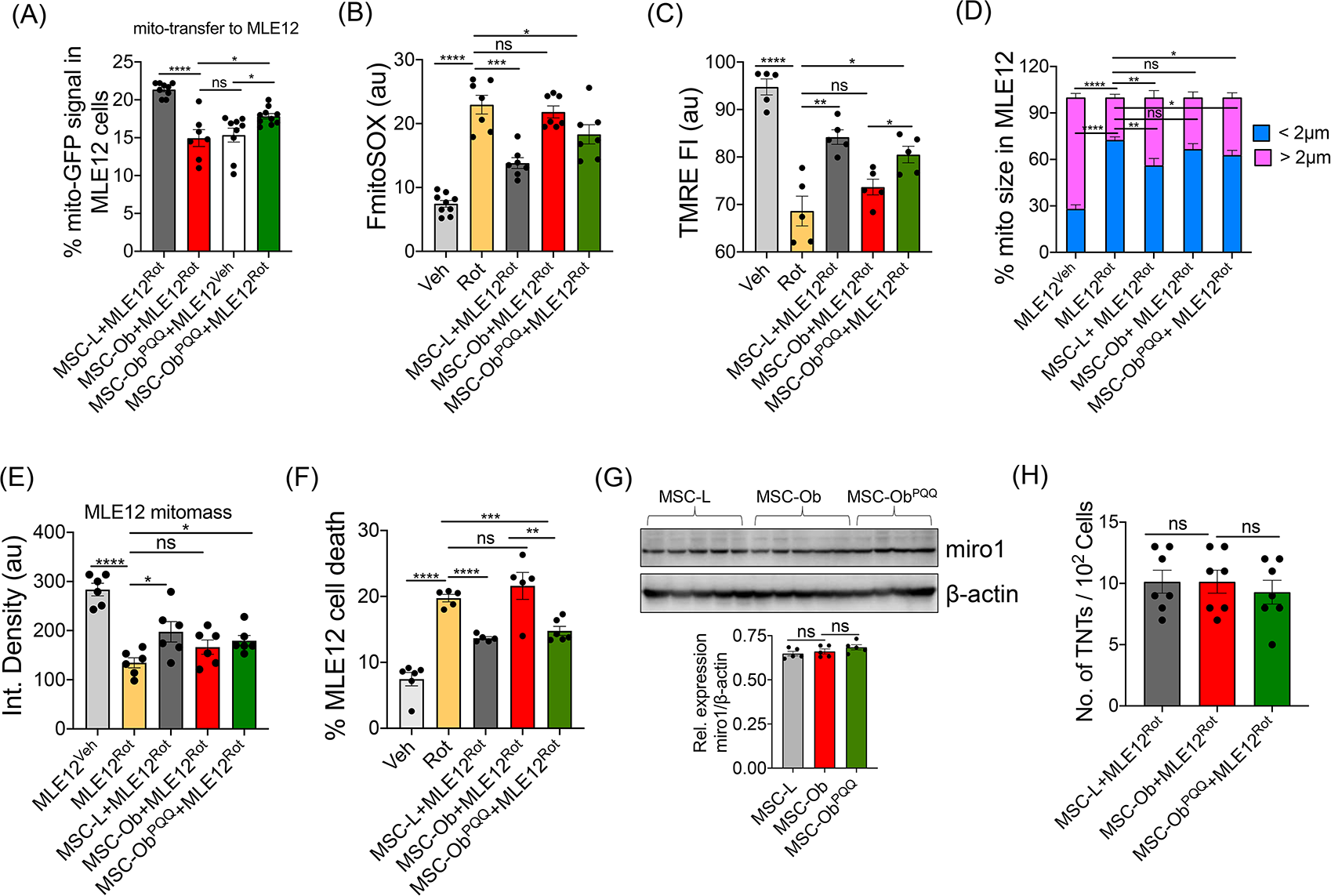
Intercellular mitochondrial transport and therapeutic potential of MSC-Ob are restored upon culturing in PQQ: **(A)** MSCs were transduced with mito-GFP and co-cultured with MLE12 which were treated with Veh or Rot and stained with CTDR. The % GFP signal was counted in by gating MLE12 cells after 24 hrs. **(B, C)** Similarly, mitoSOX and TMRE staining was done in MLE12 cells after co-culture with MSCs and represented as TMRE fluorescent intensity (FI). (**D**) CTDR stained MLE12 cells were co-cultured with mito-GFP transduced MSCs and further stained with MTR. The MTR images were taken from MLE12 cells to determine mitochondrial size distribution after 48 hrs of co-culture. (**E**) Similar to *(D)*, with integrated density representing the mitochondrial mass in MLE12 cells by specifically counting the MTR signal in these cells. (**F**) MLE-12 cells were stained with CTDR and co-cultured with untagged MSCs for 24 hrs. The cells were stained with propidium iodide (PI), and flow cytometry analysis was done. MLE12 cells were gated using CTDR, and the PI signal was calculated, which is represented as % MLE12 cell death. (**G**) Immunoblots and densitometry analysis (lower panel) of Miro1 and β-actin in cell lysates prepared from various groups of MSCs. (**H**) TNT formation between co-cultured MSCs and MLE12 cells was determined by counting the number of physically attached TNTs. The data is represented as no. of TNTs per 10^2^ cells. Mean±SEM. \*\*\*\**P* < 0.001; \*\*\**P* < 0.005; \*\**P* < 0.01; \**P* < 0.05; ns (non-significant).

### MSC-Ob^PQQ^ attenuates allergic airway inflammation in an Ova-induced mice model

To investigate the effect of PQQ treatment on the therapeutic efficacy of obese derived MSCs *in vivo*. We used our established ovalbumin-induced allergic airway inflammation (AAI) model (*8*). Before transplantation, MSC-Ob were acutely (single dose for 48 hrs; MSC-Ob^P48^) or chronically (6 doses for 10 days; MSC-Ob^PC^) treated with PQQ. Further, obese mice were fed with two different doses of PQQ (2 mg and 4 mg/kg body weight) for 15 days before harvesting MSCs. Following two sensitizations and 4-day ova-challenge, we intratracheally administered MSCs (1×10^6^). The mice were sacrificed 48 hrs post-transplantation. We observed that MSC-L and MSC-Ob^PQQ^ significantly reduced airway hyperresponsiveness (AHR) in comparison to the Ova-induced group (Ova). However, unmodulated MSC-Ob failed to attenuate AHR (Fig. S12A). Corroborating with AHR data, MSC-Ob^PQQ^ showed reduced inflammatory cell infiltration (Fig. S12B, C) and attenuated mucus secretion in the airways, whereas MSC-Ob failed to do so (Fig. S12D). Further, epithelial cell damage was significantly reduced in the animals treated with MSC-Ob^PQQ^ than MSC-Ob (Fig. S12E). Notably, MSCs harvested from obese mice which were fed PQQ, did not show significant alleviation in AHR or airway inflammation suggesting that direct in vitro modulation of MSC-Ob is a better approach.

### MSC-Ob^PQQ^ enhances intercellular mitochondrial transport and rescues airway epithelial cell damage

To investigate the effect of PQQ treatment on the therapeutic efficacy of obese derived MSCs *in vivo*. We developed a well-established allergic inflammation model (Fig. 6A) based on house dust mite (HDM) – a naturally occurring allergen and more suitable for studying allergic asthma. We chose two different time points to (1) find the effect of MSCs on intercellular mitochondrial transfer and (2) the effect on the health of airway epithelial cells and single chronic PQQ treatment (referred as MSC-Ob^PQQ^ group hereafter). To track the intercellular mitochondrial donation, we transduced MSCs with a lentiviral expressing mito-GFP. The tagged MSCs were intratracheally administered and allowed for 24 hrs before sacrifice or 48 hrs using untagged MSCs. Flow cytometry analysis and immunostaining were performed to determine the mitochondrial donation of MSCs to airway epithelial cells. The airway epithelial cells were visualized by staining with epithelial cell marker; EpCAM (for flow cytometry) and Uteroglobin (for immunofluorescence). As shown in Fig. 6B, GFP positive signal was detected in lung epithelial cells from MSC-L, while MSC-Ob showed a significantly reduced signal. Notably, MSC-Ob^PQQ^ administered animals showed restoration of the GFP signal in the airway epithelial cells (Fig. 6C, D). We further evaluated whether mitochondrial donation had any effect on the bioenergetics and mitochondrial health. To do so, we measured mtROS, ΔΨm and ATP levels in tissue lysate. In comparison to the unmodulated cells, MSC-Ob^PQQ^ showed reduced mtROS, attenuated mitochondrial depolarization and restoration in ATP levels (Fig. 6E-G). To find the effect of MSCs on epithelial cells, we obtained tissue sections from mice after 48 hrs of transplantation and performed TUNEL assay. As shown in Fig. 6H, I, a significant decrease in TUNEL positive cells was observed in MSC-Ob^PQQ^ group compared to MSC-Ob or HDM, indicating restoration of epithelial cell damage. Thus, these results demonstrate that chronic PQQ treatment enhances mitochondrial donation ability of MSC-Ob to epithelial cells *in vivo* and rescues airway epithelial cell damage.

**Fig. 6.**
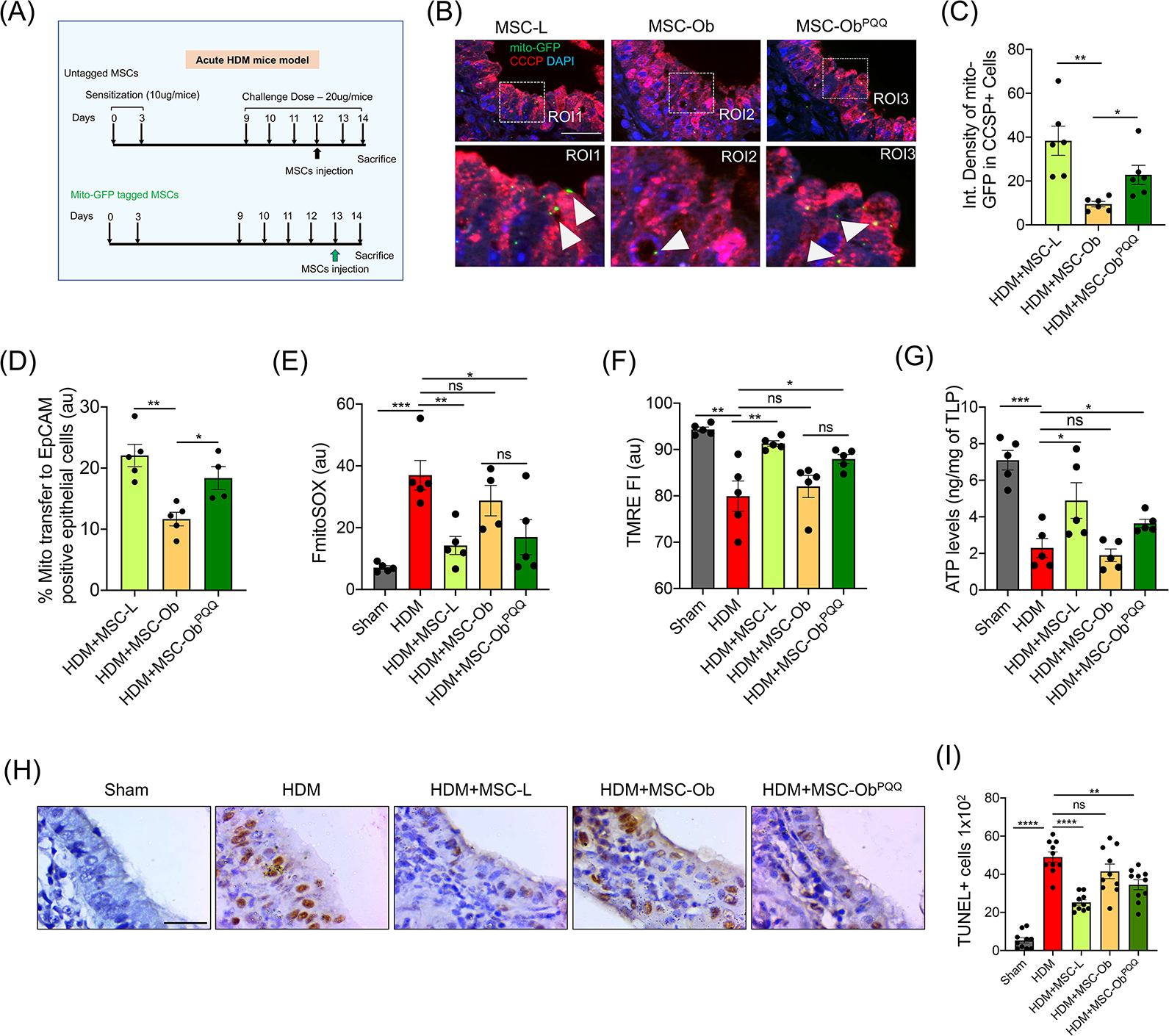
PQQ treatment restores intercellular mitochondrial transport by MSC-Ob to airway epithelial cells and enhances their therapeutic potential: (**A**) Schema showing the development of HDM-induced airway allergic inflammation model. (**B**) Representative images show Mito-GFP (green) donation by MSCs to bronchial epithelial cells stained for CCSP (RED). Mito-GFP tagged MSCs were transplanted into the mice lungs via intra-tracheal infusion, and tissue sections were prepared after 24 hrs of infusion. (**C**) Images from *(B)* were subjected to image analysis to determine the signal of mito-GFP in CCSP positive epithelial cells and represented as integrated density. (**D**) Similarly, mitochondrial donation by MSCs was calculated using flow cytometry analysis. Single cells were prepared from lung tissue and stained for EpCAM to mark epithelial cells, which were gated to calculate the GFP signal. (**E, F**) Similarly, cells were stained with mitoSOX red and TMRE, respectively, to determine the mtROS and mitochondrial membrane potential in EpCAM stained epithelial cells. (**G**) Lung tissue was obtained from different groups of mice to prepare total lung protein (TLP). The ATP levels were measured in the TLP immediately after preparation. (**H**) Representative TUNEL images from lung tissue sections prepared from animals after 48 hrs of MSC transplantation. (**I**) No. of TUNEL positive cells were calculated by counting 100 bronchial epithelial cell nuclei per section, representing cell death. Mean±SEM. \*\*\*\**P* < 0.001; \*\*\**P* < 0.005; \*\**P* < 0.01; \**P* < 0.05; ns (non-significant). Scale bars: 50 µm.

### PQQ treated MSCs attenuated AII and improved lung physiology in HDM-induced mice model

To find the effect of MSCs on airway remodeling and AHR in the HDM model, we evaluated the effect after 48 hrs of MSC transplantation. PQQ treated MSCs significantly attenuated AHR and reduced airway inflammatory cells, while MSC-Ob group showed a trend towards AHR and increased inflammatory response (Fig. 7A, B). Further, a decrease in mucus hypersecretion was observed in animals treated with MSC-Ob^PQQ^ but no significant effect was seen with MSC-Ob (Fig. 7C). To look at the effect of MSC-Ob^PQQ^ on inflammatory cells and pro-inflammatory cytokines, we performed total leucocyte count (TLC), and eosinophil cell count in BALF, and measured the inflammatory Th2 cytokine levels in the lung homogenates. A significant reduction in inflammatory cell number and Th2 cytokine release was found in mice treated with MSC-Ob^PQQ^ (Fig. 7D-H). However, MSC-Ob failed to attenuate these inflammatory responses.

**Fig. 7.**
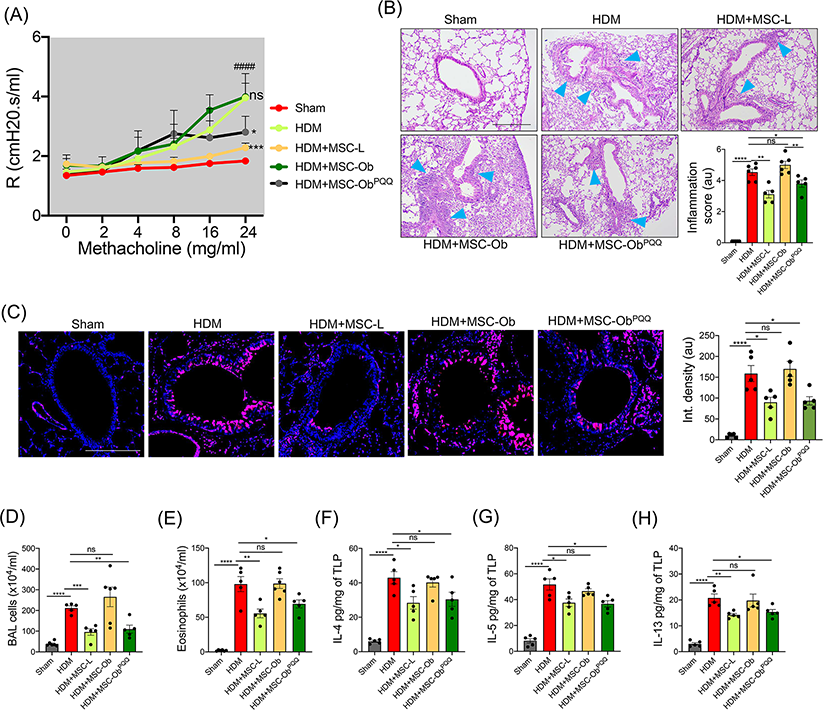
PQQ treated MSC-Ob restore airway mechanics and physiology in HDM-induced allergic asthma model: (**A**) AHR measured under various concentrations of methacholine in mice transplanted with untagged MSCs for 48 hrs before the measurements were taken. (**B**) Representative H&E images of HDM-induced allergic asthma mice transplanted with MSCs for 48 hrs. Blue arrowheads show the inflammatory cells around bronchi and blood vessels. *The lower right panel* shows inflammatory scoring representing the extent of inflammatory cell infiltration. (**C**) Representative images of tissue sections stained with PAS and pseudocolored. Pink color shows the mucus secretion, and blue shows the nuclei stained with hematoxylin. The images were subjected to image analysis to measure the PAS mucus secretion and represented as integrated density (*right panel*). (**D**) Total cell count was done in the Bronchoalveolar lavage (BAL) fluid. Similarly, eosinophil cell count was done in the BAL fluid. (**E-H**) Th2 cytokine (IL-4, IL-5, and IL-13) levels were measured in the total lung protein prepared from the lung tissue. The data is represented as a picogram of cytokines per milligram of the TLP. Mean±SEM. \*\*\*\**P* < 0.001; \*\*\**P* < 0.005; \*\**P* < 0.01; \**P* < 0.05; ns (non-significant). Scale bars: B: 200 µm; D: 100 µm.

Together with the results from the above-mentioned Ova-induced model, these results strongly suggest that MSCs derived from obese animals have compromised therapeutic efficacy, which is restored upon long-term culture of these cells in PQQ. Thus, it is imperative to consider metabolic modulation of MSCs derived from patients with metabolic syndrome before their application for therapeutic intervention.

## Discussion

In this study, we have uncovered a critical molecular pathway responsible for diminished therapeutic efficacy of MSCs derived from obese source. We found that MSC-Ob: (1) display mitochondrial dysfunction, with reduced protective intercellular mitochondrial transport; (2) show inadequate activation of mitophagic pathway; and (3) exhibit reduced cardiolipin content, which diminishes the sequestration of depolarized mitochondria into autophagosomes. These metabolic changes are together responsible for the therapeutic decline in these cells. Notably, we demonstrate that treatment with a small antioxidant molecule PQQ reverses this therapeutic deficit both in in vitro and in pre-clinical models, summarized in Fig. 8.

**Fig. 8.**
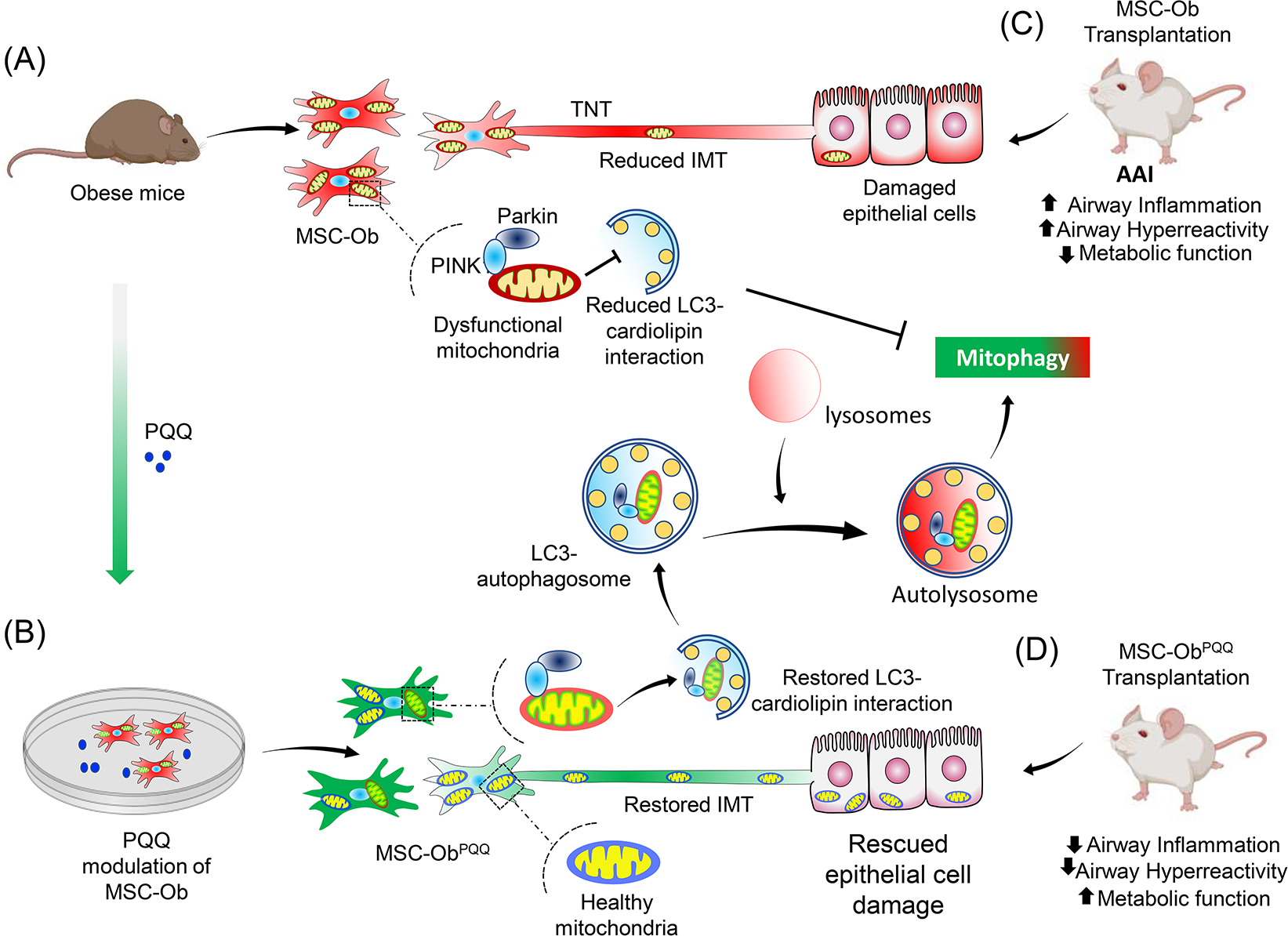
**(A)** MSCs obtained from (HFD)-induced obese mice (MSC-Ob) show underlying mitochondrial dysfunction with a concomitant decrease in cardiolipin content. These changes prevent LC3-cardiolipin interaction, thereby reducing dysfunctional mitochondria sequestration into LC3-autophagosomes and thus impaired mitophagy. The impaired mitophagy is associated with reduced intercellular mitochondrial transport (IMT) via tunneling nanotubes (TNTs) between MSC-Ob and epithelial cells in co-culture or *in vivo*. **(B)** PQQ modulation in MSC-Ob restores mitochondrial health, cardiolipin content, and thereby sequestration of depolarized into the autophagosomes to alleviate impaired mitophagy. Concomitantly, MSC-Ob shows restoration of mitochondrial health upon PQQ treatment (MSC-ObPQQ). During co-culture with epithelial cells or transplantation *in vivo* into the mice lungs, MSC-ObPQQ restores IMT and prevents epithelial cell death. (**C**) Upon transplantation in two independent allergic airway inflammatory mice models, MSC-Ob failed to rescue the airway inflammation, hyperactivity, metabolic changes in epithelial cells. (**D**) PQQ modulated MSCs restored these metabolic defects and restored lung physiology and airway remodeling parameters.

Age and dietary factors influence the metabolic state of cells with prominent changes in mitochondrial form and function (*64, 65*). MSCs derived from human adipose tissue were shown to possess mitochondria with lowered oxygen consumption rate and compromised intrinsic mitochondrial respiration parameters (*66*). We provide the first direct evidence that a high-fat diet modulates the metabolic state and sensitizes MSCs to apoptosis under culture. Using an in vitro model of IMT as a rescue assay, we demonstrate that while MSC-L restored mtROS levels, membrane potential, bioenergetics, and apoptosis of metabolically stressed MLE12 cells in coculture, MSC-Ob significantly lacked the mitochondrial donation potential and protective effect. Notably, the decline in mitochondrial donation was independent of Miro1 and changes in TNT formation. This diminished IMT was due to underlying mitochondrial dysfunction which is consistent with our previous study that chemically induced mitochondrial dysfunction diminishes IMT by MSCs (*8*).

Normally, dysfunctional mitochondria are eliminated from cells by selective autophagy (mitophagy) to escape cell death (*34*). However, MSC-Ob displayed impaired mitophagy with concomitant accumulation of dysfunctional mitochondria. Generally, the mitophagy is initiated by stabilization of PINK1 on the depolarized mitochondria which then recruits the cytosolic Parkin to ubiquitinate the OMM proteins (*50*). The ubiquitin-decorated mitochondria are then loaded to LC3-phagophores to form the autophagosome before being eventually cleared by lysosomes (*50*). Interestingly, our data shows that despite accumulation of PINK1 and Parkin on the inherently depolarized mitochondria of MSC-Ob, these cells are unable to clear the damaged mitochondria. Evaluating the downstream pathway, we found defect in the sequestration of mitochondria to LC3-containing autophagosomes as the underlying cause.

LC3 binds to mitochondria via mitophagy receptors such as FUNDC, p62, AMBRA1, BNIP3, NIX or via interaction with the ubiquitin chains associated with mitochondrial proteins (*67–69*). While we have not specifically investigated the role of all these proteins, we found that at least FUNDC and p62 may not be directly involved. Previous studies in animal models of (HFD)-induced obese mice have reported that FUNDC1 knockout aggravated obesity and insulin resistance (*70*). However, MSC-Ob did not exhibit altered expression of FUNDC1, which led us to explore other non-canonical pathways. Interestingly, we found substantial decrease in the cardiolipin content of MSC-Ob, which is reported to interact with LC3 during mitophagy (*58*). We also found significant decrease in the LC3 and cardiolipin colocalization in MSC-Ob and in human MSCs treated with FFA (**signal 1**). Based on these findings, we propose cardiolipins as putative mitophagy receptors for LC3 in MSCs and an underlying mechanism that prevents sequestration of dysfunctional mitochondrial to LC3-autophagosomes. In addition, we found a significant decrease in the induction of LC3-autophagosomes in MSC-Ob at basal and chemically depolarized conditions (**signal 2**). Contrary to the report that over-fed state reduces autophagy-related gene expression (*56*), we could not find any significant changes in the expression of key autophagy proteins such as Beclin1, Atg5, and Atg7 or gene expression by performing the autophagy pathway specific expression analysis; we presume that other regulatory mechanisms may exist (*46*). Recent studies have reported that cells deficient in LC3-dependent autophagosome formation use alternative pathways to clear damaged mitochondria, such as Rab GTPase mediated endosomal-lysosomal pathway or mitochondrial-derived vesicles (MDV)-lysosomal pathways (*42, 53, 54*). We evaluated whether MSC-Ob utilize the former pathway. While we found an increase in late endosomal/lysosomal content in MSC-Ob, surprisingly the lysosomal content did not correlate with mitochondrial clearance, thus negating the possibility of this alternate mitophagy pathways in our system. To eliminate the role of defective lysosomal function, we used a control autophagy assay based on p62 sequestration to lysosomes. We found recruitment of autophagy protein p62 to lysosomes suggesting existence of a functional lysosomal pathway. Thus, we propose a model wherein a cumulative effect (signal 1 and signal 2) of decrease in sequestration of dysfunctional mitochondria to autophagosomes and reduced induction of these structures leads to impaired mitophagy and decrease in IMT in MSC-Ob. In some instances, as a survival mechanism, MSCs outsource mitophagy to remove their partly depolarized mitochondria (*42*), or by shedding off damaged regions of mitochondria as MDVs (*71*). Although we did not evaluate these alternate mitophagy pathways, our results encourage future studies to explore these mechanisms.

As it became evident that MSC-Ob has impaired mitophagy, we sought ways to restore the mitochondrial function and induce mitophagy which may subsequently restore IMT. A small-molecule screen led to PQQ identification, which alleviated mitochondrial dysfunction and restored IMT. Notably, PQQ enhances cardiolipin content LC3-dependent autophagosome formation with concomitant induction of mitophagy to eliminate depolarized mitochondria. These results point toward a new role of PQQ in inducing mitophagy in MSCs, besides regulating mitochondrial biogenesis and mtROS, as shown by others (*60, 72*). Our findings here are consistent with a recent report showing that PQQ induces autophagy in human microglia by regulating LC3 maturation and Atg5 expression, independent of its effect on mitochondrial biogenesis (*61*). Further, in neuroblastoma cells and mice models, PQQ induces AMPK1 expression, which is a crucial regulator of autophagosome formation (*62*). Thus, we propose that PQQ may have a multifactorial role (including in mitochondrial biogenesis), with a more pronounced effect on restoring mitochondrial health by regulating mitophagy.

To determine the therapeutic efficacy of MSC-Ob and check whether modulation of these cells with PQQ will be clinically relevant, we tested these cells in two different allergic airway inflammation models. In line with our *in vitro* findings, MSC-Ob^PQQ^ showed enhanced IMT to lung epithelial cells, restoring their bioenergetics, mitochondrial membrane potential, mtROS, and epithelial cell damage. Moreover, MSC-Ob^PQQ^ alleviated Th2 cytokine levels and inflammatory cell infiltration into the lungs, while MSC-Ob failed to do so. These results are consistent with our previous study that MSCs with chemically induced mitochondrial dysfunction fail to restore epithelial cell damage and airway remodeling (*8*). Other studies have also reported the poor therapeutic outcome of MSCs with chemically induced mitochondrial dysfunction (*11*). Thus, the metabolic activity of MSCs is emerging as integral to their therapeutic efficacy (*73–75*). In line with our work, a recent study has shown that reduced heparan sulfates in white adipose tissue macrophages reduces their ability to receive the mitochondria from adipocytes in an HFD model (*76*). Taken these reports along with our own study into consideration, we propose that obesity has a long-lasting impact on MSCs, which besides reducing their mitochondrial donation capacity, may also hamper their immunomodulatory activity (*77*). Reportedly, oxidative stress and mitochondrial dysfunction impair the immunomodulatory activity of MSCs (*78, 79*).

Our findings have far-reaching clinical applications, particularly when autologous MSC transplantation is required. Autologous MSCs are preferable over allogenic sources for their intrinsic property to alleviate graft vs. host disease (GvHD). While alternate sources such as induced pluripotent cell (iPSC)-derived MSCs (*80*) are being clinically evaluated as universal cells, their clinical efficacy and long-term safety remains unknown. At present, more than 1000 clinical trials are underway to explore the beneficial effect of MSCs (autologous and allogeneic) for the treatment of a wide range of human diseases, including complex lung disorders and obesity (*2, 81*). Thus, it is imperative to evaluate the outcome of these studies by considering the source of MSCs. Based on our findings, we strongly recommend evaluating the impact of tissue source of MSCs and suggest that the cells derived from diseased patients should undergo rational modulation for clinical application.

## Material and Methods

### Study Approval

All animal experiments were approved by Institutional Animal Ethics Committee, Institute of Genomics and Integrative Biology (IGIB/IAEC/4/29/March 2018). Human bone marrow samples were obtained from healthy individuals with no history of Metabolic syndrome and Obesity at Apollo Indraprastha Hospital, New Delhi. The patient consent was taken before collecting the samples and use in the study.

### Cell Lines, Isolation and characterization of mouse and human mesenchymal stem cells

MLE12 (mouse lung epithelial cells), were purchased from ATCC and cultured as per recommendation. Mouse mesenchymal stem cells (MSCs) were isolated from the bone marrow of C57 BL/6 mice using previously described protocol (*82*). The marrow-derived cells were cultured in Mesencult (Stem Cell Technologies, Canada), which is a defined selective growth medium for mouse MSCs. The primary cells were cultured in a plating density of 10^6^ cells/ml in T25 culture flasks, and all experiments were performed in third passages.

Mouse MSCs were characterized using Sca-1-PE (eBioscience, USA), CD-44-PE (eBioscience, USA), CD11b-FITC (eBioscience, USA), Isotype-Rat IgG2ak-PE (eBioscience, USA) and Isotype-Rat IgG2bk-FITC (eBioscience, USA) using manufacturer’s protocol (BD biosciences, USA). 10,000 cells were acquired using BD FACS Melody and data were analyzed using FlowJo software.

Human MSCs were isolated from the bone marrow of healthy individual as previously described (*83*). Briefly, 1ml of bone marrow was incubated in one well of 6-well plate with 3 ml StemPro™ MSC serum-free medium (Gibco, Thermo Scientific). Media was changed after every two days until it reached to confluency. Cells were characterized in third passage using following antibodies: CD29—FITC (eBioscience, USA), CD73-PE (BD, USA), HLA-Class II-FITC (BD, USA), CD34/45-PE/45 (BD, USA), CD44-PE (eBioscience, USA), Mouse IgG1k-PE (eBioscience, USA), Mouse IgG2bk-PE (eBioscience, USA) and Rat IgG2bk-FITC (eBioscience, USA). For each marker analysis, around 10,000 cells were acquired using flow cytometer (BD Melody).

### Development of high-fat diet-induced obese mice model

High-fat diet-induced obesity model was developed using our previously described method (*47*). Briefly, four to six-week-old C57BL/6 male mice were divided in two groups and were labelled according to the diet provided as control (Chow diet) and high-fat diet (HFD) (Research Diets, Inc.). All the mice were housed in IVC cage with enrichment facilities. Every week, weight estimation was done to record the weight gain. Animals were sacrificed using the combination of xylazine and thiopentone sodium as per body weight.

### Development of HDM and Ova-induced allergic asthma mice models

Acute mice model of asthma using Ovalbumin (OVA) (Sigma) and House dust mite (HDM) (Greer Laboratories) were developed as previously described (*8*). Briefly, Ova group of mice were sensitized with a mixture of 50μg OVA and 4 mg alum (Vehicle) dissolved in 1X PBS and control group (Sham) were sensitized with 4 mg alum dissolved in 1X PBS on 0, 7^th^ and 14^th^ day. Later mice were challenged with vehicle and OVA from 21^st^ to 27^th^ day. MSCs were administered to the treatment group by intra-tracheal route 48 hrs prior to sacrifice. HDM group of mice were sensitized with HDM (10μg/mice) and Sham with 1XPBS on day 0 & day 3. Challenge of HDM and vehicle were given to mice from day 9^th^ to 14^th^. Un-transduced MSCs were intratracheally administered to treatment group (HDM challenged) 48 hrs prior sacrifice whereas, Mito-GFP transduced MSCs were transplanted 24 hrs prior sacrifice.

### Drug treatments

For initial in vitro screening to find the candidate drug molecule which enhances mitophagy by restoring depolarized mitochondria colocalisation with the LC3-autophagosomes, we used the following drugs: Pyrroloquinoline quinone (PQQ; 30µM); nicotinamide mononucleotide (NAM; 1mM); Urolithin A (UA; 50µM); Mito-Tempo (Mito-T; 100 µM); resveratrol (RSV: 1 µM); 3-methyladenine (3-MA; 5mM); N-acetyl cysteine (NAC; 1mM). After 48 hrs of incubation, the cells were fixed and stained for LC3 and Tom20. Images were taken and subjected to analysis using Image J to calculate Integrated density and Mander’s coefficient.

For all other in vitro studies MSCs were treated with PQQ (30 μM) at different time intervals. Cells were treated with 6 doses of PQQ (chronic) and media containing PQQ was changed after every 48 hrs. For *in vivo* studies cells were treated with a single dose of PQQ for 48 hrs (MSC-Ob^P48^) or 6 doses of PQQ (MSC-Ob^PQQ^) with media change after every 48 hrs. Further, HFD mice were fed 2 mg or 4 mg/kg body weight of PQQ by oral gavage in 20 µl of water. PQQ was given daily for a period of 15 days after which the mice were sacrificed and MSCs were harvested.

Human MSCs were treated with a mixture of FFA such as Palmitic acid, Stearic acid, and Oleic acid, ratio of 1:1:1). Cells were treated with two different concentrations of FFA (750 mM and 1 mM) for 24 hrs (*59*). The PQQ treatment (30μM) was done 2 hrs prior to FFA treatment.

### Measurement of blood glucose and biochemical analysis

Blood glucose was measured using Accu-chek active test strips as earlier described (*47*). Biochemical parameters such as triglycerides and cholesterol level were measured in the blood serum by quantitation kits (BIOVISION) as previously described (*47*).

### Cellular Senescence Assay

MSCs were seeded in 24 well plates and staining for cellular senescence was performed using manufacturer’s protocol (Sigma). Briefly, cells were washed with 1X PBS and fixed with fixative buffer for 7 mins at room temperature. Cells were stained with SA-β-gal staining solution at 37 °C for 4 hrs. Senescent cells were stained blue and counted using phase contrast microscope at 10X magnification. The percentage was calculated from five different view field of each sample in four independent experiments.

### Annexin V and propidium cell death assay

The cells were dissociated with 0.125% trypsin-EDTA (Sigma) and then stained with using Dead cell apoptosis kit (Invitrogen) with Annexin V FITC and propidium iodide using manufacturer’s protocol. 10,000 cells were acquired using flow cytometry (BD Melody).

### Live and Dead Assay

The live and dead assay was performed using the LIVE/DEAD™ Viability/Cytotoxicity Kit (Invitrogen) according to the manufacturer’s instructions. Briefly, the medium was changed to phenol red-free DMEM with green fluorescent calcein-AM to indicate intracellular esterase activity (live cells) and red-fluorescent ethidium homodimer-1 which indicates loss of plasma membrane integrity (dead cells). Cells were incubated at 37 °C for 30 mins and were analyzed under FLoid™ Cell Imaging Station.

### Lentiviral Production

Lentiviral particle packaging was performed using previously described method (*84*). Briefly, Plasmid encoding for mitochondria specific protein/ autophagosome specific protein is transfected in HEK293T cells along with packaging vector (pDR8.2; Addgene #8455) and envelope encoding protein (VSVG; Addgene # 8454). Lentiviral particles were collected after 48 hrs of transfection and were used for transducing target cell lines.

### Live cell imaging

MSCs were transduced with lentiviral particles having mitochondria targeting protein and GFP in downstream (Addgene) and lysosome were stained with lysotracker Deep-Red (Invitrogen, USA) followed by 3 times wash with 1X PBS. Imaging was done before and after treating with FCCP (10 uM) for 2 hrs using Nikon confocal Ti2E at 60x magnification. Image analysis was done using Nikon Elements software and Image J.

Quantitation of mitochondrial size in mouse and human MSCs were done by staining with Mito-tracker Red (Invitrogen/Thermo Fisher Scientific) using manufacturer’s protocol. In brief, cells were stained with mito-tracker red for 15 mins at 37 °C, followed by 3 wash with 1X PBS. Imaging was done using Nikon confocal Ti2E at 60x magnification and analysis was done using Image J.

### Immunofluorescence

Cells were allowed to adhere to glass coverslip before being fixed with 4% paraformaldehyde (ThermoFisher Scientific) in 1X PBS for 15minutes at room temperature. Permeabilization and blocking was done in a buffer containing 0.1% Triton X-100, 5% goat serum in 1X PBS for 1 hr at RT. Primary antibodies, shown in Table 1, incubation was done in the buffer containing 0.01% Triton X-100 and 2% goat serum in 1X PBS overnight at 4 °C. Secondary antibody incubation was done in the same buffer using fluorophore tagged secondary antibodies for 1 hr at RT. Cells were washed between steps using 1X PBS for 5 minutes each at RT. The coverslip was then mounted on frosted slides (Corning) using DAPI mountant and allowed to dry before being sealed using colorless nail polish. TNT were visualized by staining F actin with phalloidin 594 (Invitrogen).

**Table 1:** Key resources table.

| **Primary Antibodies** |  |  |  |
| --- | --- | --- | --- |
| **S. No.** | **Name** | **Company** | **Catalogue No.** |
| 1 | Anti-Mitofusin 2 | Abcam | ab56889 |
| 2 | Recombinant Anti-DRP1 | Abcam | ab184247 |
| 3 | Anti-PGC1 alpha | Abcam | ab54481 |
| 4 | Anti-MIRO1 | Abcam | ab83779 |
| 5 | Anti-LC3B | Abcam | ab48394 |
| 6 | Anti-TOMM20 | Abcam | ab56783 |
| 7 | Anti-LAMP1 | Abcam | ab25245 |
| 8 | Recombinant Anti-Beclin 1 | Abcam | ab207612 |
| 9 | Anti-VDAC1/Porin | Abcam | ab15895 |
| 10 | Anti-PGC1 alpha | Abcam | ab106814 |
| 11 | Anti-PINK1 | Abcam | ab75487 |
| 12 | Anti-COX IV | Abcam | ab14744 |
| 13 | Anti-FUNDC1 | Abcam | ab74834 |
| 14 | Anti-DRP1 | Abcam | ab140494 |
| 15 | Anti-SQSTM1 / p62 | Abcam | ab56416 |
| 16 | Anti-Uteroglobin | Abcam | ab40873 |
| 17 | Recombinant Anti-EpCAM | Abcam | ab32392 |
| 18 | Anti-alpha Tubulin | Abcam | ab7291 |
| 19 | Anti-Mitofusin 1 | Abcam | ab57602 |
| 20 | Anti-LAMP1 | Abcam | ab24170 |
| 21 | Anti-Cytochrome C | Abcam | ab90529 |
| 22 | Anti-Parkin | Cell Signaling Technology | 2132S |
| 23 | Anti-Phospho Beclin1 | Cell Signaling Technology | 13825 |
| 24 | Anti-COX IV | Cell Signaling Technology | 4844 |
| 25 | Anti-Atg5 | Cell Signaling Technology | 2630 |
| 26 | Anti-Atg7 (D12B11) Rabbit mAb | Cell Signaling Technology | 8558 |
| 27 | Anti-PINK1 Rabbit pAb | ABclonal | A11435 |
| 28 | Anti-Parkin Rabbit pAb | ABclonal | A0968 |
| 29 | Anti-β-Actin antibody, Mouse monoclonal | Sigma | A1978-200UL |
| 30 | CD326 (EpCAM) Monoclonal Antibody (G8.8), PE | eBioscience | 12-5791-82 |
| 31 | Ly-6A/E (Sca-1) Antibody, PE conjugated (Mouse) | eBioscience | 12-5981-82 |
| 32 | CD105 Antibody PE conjugated (Mouse) | eBioscience | 12-1051-82 |
| 33 | CD11b Antibody FITC conjugated (Mouse) | eBioscience | 11-0112-82 |
| 34 | CD44 Monoclonal Antibody (IM7), PE | eBioscience | 12-0441-82 |
| 35 | CD29 (Integrin beta 1) Monoclonal Antibody (TS2/16) | eBioscience | 14-0299 |
| 36 | Rat IgG2a kappa Isotype Control (eBR2a), PE | eBioscience | 12-4321-80 |
| 37 | Rat IgG2b kappa Isotype Control (eB149/10H5), FITC | eBioscience | 11-4031-82 |
| 38 | Mouse IgG2b kappa Isotype Control (eBMG2b), PE | eBioscience | 12-4732-41 |
| 39 | PE Mouse Anti-Human CD73 | BD Biosciences | 550257 |
| 40 | PE-Cy<sup>TM</sup>5 Mouse Anti-Human CD90 | BD Biosciences | 555597 |
| 41 | FITC Mouse Anti-Human HLA-DR | BD Biosciences | 555560 |
| 42 | Anti-Human CD45 FITC/CD34 PE | BD Biosciences | 341071 |
| 43 | PE Mouse IgG1, κ Isotype Control | BD Biosciences | 554080 |
| **Secondary Antibodies** |  |  |  |
| 1 | Donkey anti-Rabbit IgG (H+L) Highly Cross-Adsorbed Secondary Antibody, Alexa Fluor 488 | Invitrogen | A-21206 |
| 2 | Donkey anti-Goat IgG (H+L) Cross-Adsorbed Secondary Antibody, Alexa Fluor 488 | Invitrogen | A11055 |
| 3 | Goat anti-Rabbit IgG (H+L) Highly Cross-Adsorbed Secondary Antibody, Alexa Fluor 594 | Invitrogen | a11037 |
| 4 | Donkey anti-Rabbit IgG (H+L) Highly Cross-Adsorbed Secondary Antibody, Alexa Fluor 647 | Invitrogen | a31573 |
| 5 | Donkey anti-Mouse IgG (H+L) Highly Cross-Adsorbed Secondary Antibody, Alexa Fluor 488 | Invitrogen | a21202 |
| 6 | Donkey anti-Mouse IgG (H+L) Highly Cross-Adsorbed Secondary Antibody, Alexa Fluor 546 | Invitrogen | a10036 |
| 7 | Goat anti-Rat IgG (H+L) Cross-Adsorbed Secondary Antibody, Alexa Fluor 647 | Invitrogen | A21247 |
| 8 | Rabbit anti-mouse IgG - HRP | GeNei<sup>TM</sup> | 1140580011730 |
| 9 | Rabbit anti-goat IgG – HRP | GeNei<sup>TM</sup> | 1140480011730 |
| 10 | HRP-AffiniPure Goat Anti-Rabbit IgG (H+L) Secondary Antibody | Jackson ImmunoResearch Inc. | 111-035-003 |
| **Serum** |  |  |  |
| 1 | NORMAL GOAT SERUM | Jackson | 005-000-121 |
| 2 | NORMAL RABBIT SERUM | Jackson | 011-000-120 |
| **Drugs** |  |  |  |
| 1 | Urolithin A <sup>(1)</sup> | Sigma | SML1791 |
| 2 | Resveratrol <sup>(2)</sup> | Sigma | R5010 |
| 3 | Mito-Tempo<sup>(3)</sup> | Santa Cruz | SC-221945 |
| 4 | Pyrroloquinoline quinone <sup>(4)</sup> | Sigma | D7783 |
| 5 | Nicotinamide mononucleotide <sup>(5)</sup> | Sigma | N3501 |
| 6 | 3-methyladenine <sup>(6)</sup> | Sigma | M9281 |
| 7 | N-acetyl cysteine <sup>(7)</sup> | Sigma | A7250 |
| 8 | Carbonyl cyanide 4-(trifluoromethoxy)phenylhydrazone | Sigma | C2920 |
| 9 | Oligomycin A | Sigma | 75351 |
| 10 | Antimycin A | Sigma | A8674 |
| 11 | Rotenone | Sigma | R8875 |
| 12 | Sodium Palmitate | Sigma | P9767 |
| 13 | Stearic Acid | Sigma | S4751 |
| 14 | Oleic acid | Sigma | O1383 |
| **Plasmids** |  |  |  |
| 1 | pLYS1-FLAG-MitoGFP-HA | Addgene | 50057 |
| 2 | pCMV-VSV-G | Addgene | 8454 |
| 3 | pCMV-dR8.2 dvpr | Addgene | 8455 |
| 4 | pLVX-LC3-YFP | Addgene | 99571 |
| **Cell staining dyes** |  |  |  |
| 1 | TMRE | Invitrogen | T669 |
| 2 | MitoSOX Red | Invitrogen | M36008 |
| 3 | Mitotracker Red | Invitrogen | M7513 |
| 4 | Mitotracker Green | Invitrogen | M7514 |
| 5 | Lysotracker green | Invitrogen | L7526 |
| 6 | Lysotracker deep red | Invitrogen | L12492 |
| 7 | Propidium Iodide | Invitrogen | V13242 |
| 8 | Hoechst 33342 | Invitrogen | H21492 |
| 9 | CellTrace™ CFSE Cell Proliferation Kit | Invitrogen | C34554 |
| 10 | CellTracker™ Green | Invitrogen | C2925 |
| 11 | CellTracker™ Deep Red Dye | Invitrogen | C34565 |
| 12 | DAPI | Invitrogen | P36966 |
| 13 | Alexa Fluor™ 594 Phalloidin | Invitrogen | A12381 |
| 14 | Nonyl Acridine Orange (NAO) | Invitrogen | A1372 |
| 8 | Hoest 3344 | Invitrogen | H21492 |
| 9 | CFSE | Invitrogen | C34554 |
| 10 | Cell tracker green | Invitrogen | C2925 |
| 11 | DAPI | Invitrogen | P36966 |
| 12 | Phalloidin 594 | Invitrogen | A12381 |
| **Kits** |  |  |  |
| 1 | Senescence Cells Histochemical Staining Kit | Sigma | CS0030 |
| 2 | Mitochondria Isolation Kit | Sigma | MITOISO2-1KT |
| 3 | ELISA MAX™ Deluxe Set Mouse IL-4 | BioLegend | 431104 |
| 4 | ELISA MAX™ Deluxe Set Mouse IL-5 | BioLegend | 431204 |
| 5 | Live Dead assay kit | Invitrogen | L3224 |
| 6 | Dead cell apoptosis kit | Invitrogen | V13242 |
| 7 | DeadEnd™ Colorimetric TUNEL System | Promega | G7130 |
| 8 | ATP assay kit | Abcam | ab83355 |
| 9 | Mouse IL-13 DuoSet ELISA | R&D Systems | DY413 |
| **Chemicals** |  |  |  |
| 1 | Sodium Pyruvate 100mM | Sigma | S8636-100ML |
| 2 | Sodium Azide | Sigma | S8032-25G |
| 3 | (+)-Sodium L-Ascorbate | Sigma | A4034-100G |
| 4 | L-(-)-Malic Acid | Sigma | M6413-25G |
| 5 | TMPD | Sigma | T7394-5G |
| 6 | ADP | Sigma | A5285-1G |
| 7 | Succinic acid | Sigma | S9512-100G |
| 8 | Palmitoyl-L-carnitine chloride | sigma | P1645 |
| 9 | TMB liquid substrate system for ELISA | Sigma | T0440-1L |
| 10 | Hydrogen peroxide–Urea adduct | Sigma | U8879-100<br>TAB |
| 11 | Acetyl-β-methylcholine chloride | Sigma | A2251-25G |
| 12 | Hydrogen peroxide solution | Sigma | 323381-500ML |
| 13 | Bicinchoninic Acid (BCA) | Sigma | B9643-1L |
| 14 | Water: Molecular Biology Reagent | Sigma | W4502-6X1L |
| 15 | Sodium dodecyl sulfate (SDS) | Sigma | L3771-500G |
| 16 | D-Glucose | Sigma | G8270-100G |
| 17 | Triton X 100 | Sigma | T9284 |
| 18 | Hexadimethrine bromide | Sigma | H9268-10G |
| 19 | Bovine Serum Albumin Fatty acid free (BSA) | Sigma | A7511 |
| 20 | Insulin human | Sigma | I2643 |
| 21 | Protease Inhibitor Cocktail | Sigma | p8340 |
| 22 | Albumin from chicken egg white | Sigma | A5503 |
| 23 | Pierce™ 16% Formaldehyde | Invitrogen | 28908 |
| 24 | RNA later | invitrogen | AM7021 |
| 25 | cDNA synthesis kit | Invitrogen | 4368814 |
| 26 | LTX | Invitrogen | A12621 |
| 27 | SuperSignal™ West Femto Maximum Sensitivity Substrate | Invitrogen | 34095 |
| 28 | SuperSignal™ West Pico PLUS Chemiluminescent Substrate | Invitrogen | 34580 |
| 29 | L-Glutamine (200 mM) | Invitrogen | 25030081 |
| 30 | HBSS | Invitrogen | 14025092 |
| 31 | PageRuler™ Prestained Protein Ladder | Invitrogen | 26616 |
| 32 | Ampicillin sodium salt | HiMedia | CMS645-5G |
| 33 | Kanamycin sulphate | HiMedia | MB105-1G |
| 34 | Phosphate Buffered Saline Tablets | Gibco | 18912014 |
| 35 | StemPro™ MSC SFM | Gibco | A1033201 |
| 36 | KAPA SYBER FAST QPCR MM | KAPA | kk4601 |
| 37 | MesenCult Expansion Kit (Mouse) | Stem Cell Technologies | #05513 |
| 38 | Bovine Serum Albumin (BSA) | Bio World | 22070004-5 |
| 39 | RNeasy Mini Kit (250) | Qiagen | 74106 |
| 40 | House Dust Mite Extract<br>Dermatophagoides pteronyssinus, 2.5 ml | Greer Laboratories | XPB70D3A2.5 |
| 41 | Rodent purified diet W/60% energy from Fat diet | Research Diets, Inc. | D12492 |

Human and mouse MSCs were stained with NAO (Invitrogen) followed by three times wash with 1X PBS and fixed with 4% PFA. After fixation, cells were washed 3 times with 1X PBS, blocked in blocking buffer and primary antibody incubation for overnight at 4 °C followed by secondary antibody incubation for 1 hr. Mounted using DAPI and images were obtained on a Nikon confocal Ti2E and were analyzed using Nikon Elements software and Image J.

Immunofluorescence of Mito-GFP presence in MSC transplanted mice lung tissue was done as previously described (*8*). Briefly, tissue sections were deparaffinized following the antigen retrieval. Permeabilization was done in a buffer containing 0.1% Triton X-100 for 30mins followed by blocking with blocking buffer (5% sera in 0.01% Triton X-100) at RT for 1 hr. Primary antibody incubation was done in same buffer for overnight at 4 °C followed by 3 times wash with 1 X PBS. Secondary antibody incubation was done in the same buffer for 1 hr at RT followed by 3 times wash with 1X PBS. Nuclear staining was done using DAPI and mounted with DPX mounting solution using coverslip. The images were obtained on a Zeiss microscope integrated with Apotome 2 (ZEISS Axio Observer 7) and analyzed using Image J.

### Image analysis

The microscopy images were first processed that includes splitting of the three-channel image, selecting the green channel (labelled as mitochondria) image, followed by sharpen, despeckle, background subtraction and enhancing the local contrast of the final image. As mitochondria are filamentous tube-like structures, so a filter named as ‘tubeness’ (sigma = 0.0210) is also applied to enhance the filamentous property of the segmented mitochondria. At last, the image is converted into binary preceded by gaussian blur (sigma radius = 1.000). This image is then used to calculated the mitochondria length.

For integrated density calculation, the image was first split into three different channels. The green channel image is selected and rest are closed. During the image processing step, the background was subtracted from the image followed by binary conversion. Finally, the image was redirected to measure the integrated density parameter.

For colocalisation analysis, the ‘EzColocalization’ plugin (*85*) was used to measure the following parameters. At first step, the three-channel image was again split into different channels, out of which red and green channel image was taken to perform colocalization analysis. Mander’s overlap coefficient (or Mander’s’ coefficient) was calculated between PINK1 and Tom20; Parkin and Tom20; LC3 and Tom20 or mito-GFP; LAMP1 and Tom20; p62 and LC3; p62 and LAMP1; Tom20 and MTDR. Mander’s coefficient measures the percentage colocalisation between two channels with values ranging from 0 to 1 as represented in the Figure. 0 indicates no colocalisation and 1 indicates 100% colocalisation. Line scans measurements were done by drawing a 10 µm line on the images and the intensity profile of both the channels was calculated as described by us previously (*86*).

### Flow Cytometry

Mitochondria specific reactive oxygen species (ROS), membrane potential and mitochondrial mass was measured using MitoSOX Red (Invitrogen/Thermo Fisher Scientific), tetramethylrhodamine, ethyl ester (TMRE; Sigma-Aldrich, St. Louis, MO, USA) and Mito-tracker green (Invitrogen/Thermo Fisher Scientific), respectively using manufacturer’s protocol. Briefly, cells were stained under live conditions and washed 3 times with 1X PBS before acquisition. The data were presented as percentage-gated fluorescence shift and detected using BD Melody and BD Accuri C6 plus.

MSCs were stained with NAO (Invitrogen) using manufacturer’s instruction. Briefly, cells were stained with NAO, followed by 3 times wash with 1X PBS. Cells were dissociated and acquired using flow cytometer (BD Melody).

Quantification of mitochondria transfer from MSCs to MLE12 cells was done as previously described (*8*). Briefly, in *in vitro* MSCs was transduced with Mito-GFP lentiviral particles and MLE12 cells were stained with CTDR (Invitrogen/Thermo Fisher Scientific). Prior co-culture MLE12 cells were induced with vehicle (DMSO) or rotenone for 12 hrs and then co-cultured for 24 hrs at 37°C with 5% CO_2_. Quantification was done by gating double positive cells for CTDR and MitoGFP in FACS dot plot on FL3 window. For rescue experiment, cells were stained with propidium iodide (Invitrogen) using manufacturer’s protocol. Cells positive for Cell tracker green (CTG) were further analyzed for propidium iodide staining using flow cytometer (FACS Melody).

For *in vivo* study, the quantification of mitochondria transfer from MSCs to epithelial cells was done as described earlier (*8*). In brief, MSCs transduced with Mito-GFP lentiviral particles were delivered into the mice lungs by intratracheal route. Lung tissues were homogenized using Miltenyi tissue dissociator (Miltenyi Biotech, USA) and single cell suspension was prepared in 1X PBS supplemented with 1% BSA. Bronchial epithelial cells were stained with EpCAM (Ebioscience/Thermo Fisher Scientific) and quantification of mitochondria transferred to epithelial cells was done by gating double positive cells for EpCAM-PE and Mito-GFP in FACS dot plot on FL3 window.

### Mitochondrial Fraction preparation

Mitochondria fraction was prepared using mitochondria isolation kit (Sigma) following the manufacturer’ s protocol. Briefly, 2-3 x 10^7^ cells were washed with ice cold PBS and centrifuged at 600 x g @ 4 °C. The pellet was resuspended in 1.5ml of 1x extraction buffer A following with the incubation for 10 mins on ice. Cells were homogenized using Dounce homogenizer 10-30 strokes following with a centrifuge at 600x g for 10 mins @ 4 °C. Transfer the supernatant to another tube and centrifuge at 11000x g for 10 mins @ 4 °C. The supernatant contains the cytosolic fraction and the pellet obtained was mitochondria fraction. Washed the pellet with extraction buffer to remove cytosolic contamination from mitochondrial fraction.

### Western Blot analysis

Cells were lysed using RIPA buffer (Sigma) supplemented with protease inhibitor cocktail (Sigma). The debris were removed from the lysates by centrifugation and protein estimation was performed by Bicinchoninic Acid (BCA) assay (Sigma). Proteins were resolved by loading 10ug of protein on a 10% and 12% SDS-PAGE gel. Transfer of proteins to the polyvinylidene difluoride (PVDF) membrane was done using a wet transfer technique. Blots were probed with primary antibodies, indicated in Table 1, for overnight at 4 °C, followed by HRP-conjugated secondary antibodies for 1 hr at room temperature, indicated in Table 1. The protein bands were visualized with enhanced chemiluminescence (Invitrogen/Thermo scientific).

### Measurement of Oxygen Consumption Rate

Mitochondrial respiration in live MSCs were measured using seahorse XF24 Extracellular Flux Analyzer with Mito stress test kit according to manufacturer’s protocol (Agilent). In brief, cells were plated at 60,000 cells/ well onto XF24-well microplates the day before analysis. On the day of analysis, the cells were equilibrated in XF buffer and were kept in non-CO_2_ incubator for 60 minutes. OCR was measured after repeated cycles which includes mixing (3 minutes), the incubation (2 minutes), and measurement (3 minutes) periods. Following basal line measurements, cells were treated with 1 µM oligomycin (ATP synthase inhibitor), 2 µM FCCP (Mitochondrial OXPHOS uncoupler) and mixture of 1 µM rotenone (Complex I inhibitor) and antimycin A (Complex III), and the changes in OCR were recorded. OCR data was normalized by total cell protein and the data was expressed in pmol/min/ug.

### Gene Expression Analysis

Mouse MSCs RT^2^ Prolifer PCR array (PAMM-084ZG, Qiagen) was used to evaluate the expression of 84 specific genes related to autophagy using manufacturer’s instructions. Briefly, total RNA was isolated from 10^5^ to 10^6^ cells using the RNeasy Mini kit (Qiagen) using manufacturer’s protocol. cDNA synthesis from 500ng of the total RNA using the RT^2^ first strand kit (Qiagen). The RT^2^ Prolifer PCR array test includes various measures allowing to identify the contamination with genomic DNA, contamination with DNA during the procedure, to control the presence of reverse transcription and PCR inhibitors. After all control test, the samples were analyzed using the RT^2^ Prolifer PCR array, altogether 84 different genes simultaneously amplified in the sample. PCR array were performed in 384-well plates on a LightCycler 480 instrument (Roche Applied Science). The reaction mix of 102 ul of sample cDNA was prepared using 2x SA Biosciences RT^2^ qPCR Master Mix and 10 ul of this mixture was added into each well of the PCR array. The data was analyzed using Qiagen’s online Web analysis tool (https://dataanalysis2.qiagen.com/pcr). A more than twofold change in gene expression compared to control group was considered as the up-or downregulation of a specific gene expression.

The RT-qPCR for Miro1 and PCR for mitochondrial DNA was performed on Rotor gene Q (Qiagen). Relative gene expression of Miro1 and mitochondrial DNA copy number were evaluated using their specific forward and reverse primers as indicated in Table 2. The analysis and fold change expression were done as previously described (*87*).

**Table 2:**
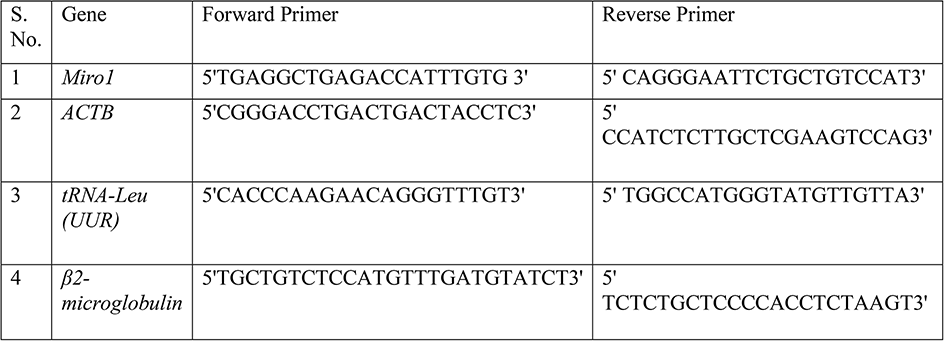
List of Primers.

### Transmission electron microscope

Examination of MSCs mitochondria using a TEM was performed as described earlier (*8*). In brief, cells were fixed overnight at 4°C in fixative containing 2.5% glutaraldehyde and 4% paraformaldehyde. After washing with 0.1M sodium cacodylate buffer to remove excess fixative, cells were embedded in 2% agar blocks. Samples were post-fixed in 2% osmium tetra oxide for 1 hr, dehydrated in graded series of ethanol (30%, 50%, 70%, and 100%) and infiltrated in epon resin and polymerized at 60°C for 72 hrs. Ultrathin sections (63nm) were cut on an Ultramicrotome (Leica EM UC7), placed on copper grids and stained with 5% uranyl acetate and 0.2% lead citrate. Sections were examined on a 200 KVA transmission electron microscope (Tecnai G2 20 twin, FEI). The condition of the cristae was analyzed manually and the mitochondria with no clear double membrane and imperfect cristae were labelled as disrupted.

### TUNEL assay

The quantification of dead cells in lung tissue was performed using an *in-situ* apoptosis detection DeadEnd Colorimetric TUNEL System (Promega) as previously described (*88*). Briefly, the tissues sections were cut into 5 µm slices and stained as per the protocol. Hematoxylin was used as a counterstain to visualize the nuclei. The images were taken with LMI microscope using 100X objective (DM-X, LMI microscopes, UK).

### ATP assay

ATP levels was measured using ATP assay kit (Colorimetric) (Abcam) following the manufacturer’s protocol in lung tissue lysate as previously described (*8*).

### Histopathology and Airway hyperresponsiveness measurement

Histopathology of lung and Airway hyperresponsiveness was measured using the protocol as previously described (*8*). The AHR was measured with the flexiVent systems (SCIREQ). The scoring of the H&E slides was done manually as described previously (*8*).

### PAS staining

The tissues sections were subjected to Periodic acid–Schiff (PAS) staining as described by us previously (*8*). Briefly, the tissues sections were deparaffinized and hydrated using decreasing concentrations of ethanol following that the slides were incubated in 0.3% periodic acid. The slides were subsequently incubated in Schiff’s reagent (Sigma) and counterstained with hematoxylin and mounted with DPX mountant (Sigma) and images were taken with LMI microscope using 20X objective (DM-X, LMI microscopes, UK). For better visualisation of the signal, the images were processed with Image J software and the images were split into three channels (red, green and blue). The red channel represents mucus secretion and the blue channels which represents nuclei (hematoxylin) were merged. The intensity of the images was calculated as described in the image analysis section and represented as integrated density.

### ELISA assay

ELISA assay of different cytokines IL-4 (BioLegend), IL-5 (BioLegend) and IL-13 (R&D Systems) was done in Lung lysate using manufacturer’s protocol. Briefly, 10 µg of the total cell lysate was used for the measurement. The readings were obtained by the spectrophotometer (Multiskan SkyHigh Microplate Spectrophotometer, Thermofischer scientific). The calculations were done as described by us previously (*8*).

### Statistical analysis

For statistical analysis, non-parametric t test was used to compare various groups. The analysis was done using Prism version 8.0 (GraphPad Software). The bar graphs were plotted in the Prism with respective P values. The data is expressed as mean±SEM and the values indicate ****P<0.001; ***P<0.005; **P<0.01; *P<0.05, which were considered as significant. The analysis was done in a minimum of three biological replicates or else as mentioned in the Fig. legends.

## Acknowledgements

We thank the lab members of Dr. Anurag Agrawal, Dr. Soumya Sinha Roy and Dr. Tanveer Ahmad for helpful discussions and suggestions during the preparation of the manuscript. We thank Dr. Mohan C Joshi (MCARS, JMI) for providing access to the microscopy facility and Dr. Naveen Kumar Bhatraju (CSIR-IGIB) for assistance with Seahorse assays. We also thank Dr. Sheetal Gandotra (CSIR-IGIB) and Stanzin Dawa (CSIR-IGIB) for assisting with lentiviral particle generation. This work was supported by the Council of Scientific and Industrial Research (CSIR), India through grant (MLP2008). Dr. Tanveer Ahmad is supported by the UGC early career grant (F.4/2018/FRP-Start-up-grant) and SERB Core research grant (CRG/2020/002294). Shakti Sagar, Imam Faizan and Nisha Chaudhary acknowledge ICMR for the Senior Research Fellowship. We thank the TEM facility of CSIR-IGIB for electron microscopy studies and Mehwish Nafiz (MCARS, JMI) for assisting in PAS staining and TUNEL assay. We also thanks Mayank Garg for assisting with *in vivo* experiments and Vandana Singh for providing free fatty acid.

## Author contribution

**S. S.** Designed and performed the experiments, compiled and analyzed the data, and assisted in writing the manuscript. **M. F. I.** Performed mitochondrial ROS assay and assisted in immunofluorescence experiments and *in vivo* studies. **N. C.** Performed image analysis, mitochondrial size analysis and co-localization studies. **A. G.** Assisted *in vivo* studies, biochemical analysis and lung function measurement. **K. S.** assisted in flow cytometry and *in vivo* studies. **I. A.** Performed cytokine assays. **V. P. S.** Assisted with development of HFD and provided the reagents for obesity characterization. **G. K.** Collected the bone marrow and assisted in human stem cell isolation. **U. M.** assisted with inflammation scoring and development of HDM mice model. **A. A.** Conceptualized the idea, supervised the study, provided the resources and assisted in manuscript writing. **T. A.** Conceptualized the idea, designed the experiments, analyzed the data, provided the resources, and wrote the manuscript. **S. S. R.** Conceptualized the idea, provided the resources, assisted in manuscript writing and supervised the overall study.

## Supplementary Figure Legends

**Figure S1.**
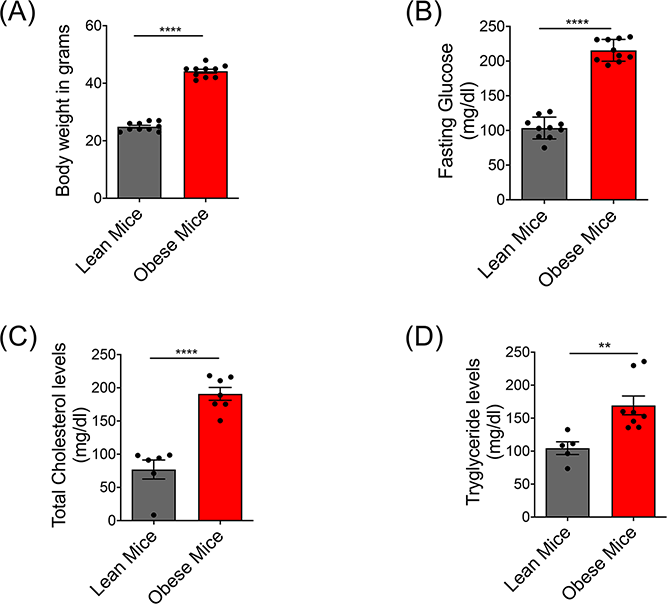
Characterization of obesity features in HFD mice model: **(A)** Body weight of the mice fed regular diet (lean mice) or high-fat diets (obese mice) was measured after 16 weeks. The weight is presented in grams/mice. (**B**) Fasting glucose was measured in the blood of lean and obese mice. (**C, D**) Similarly, total cholesterol and triglyceride content was measured in the serum. Data is presented as Mean±SEM. \*\*\*\**P* < 0.001; \*\**P* < 0.01.

**Figure S2.**
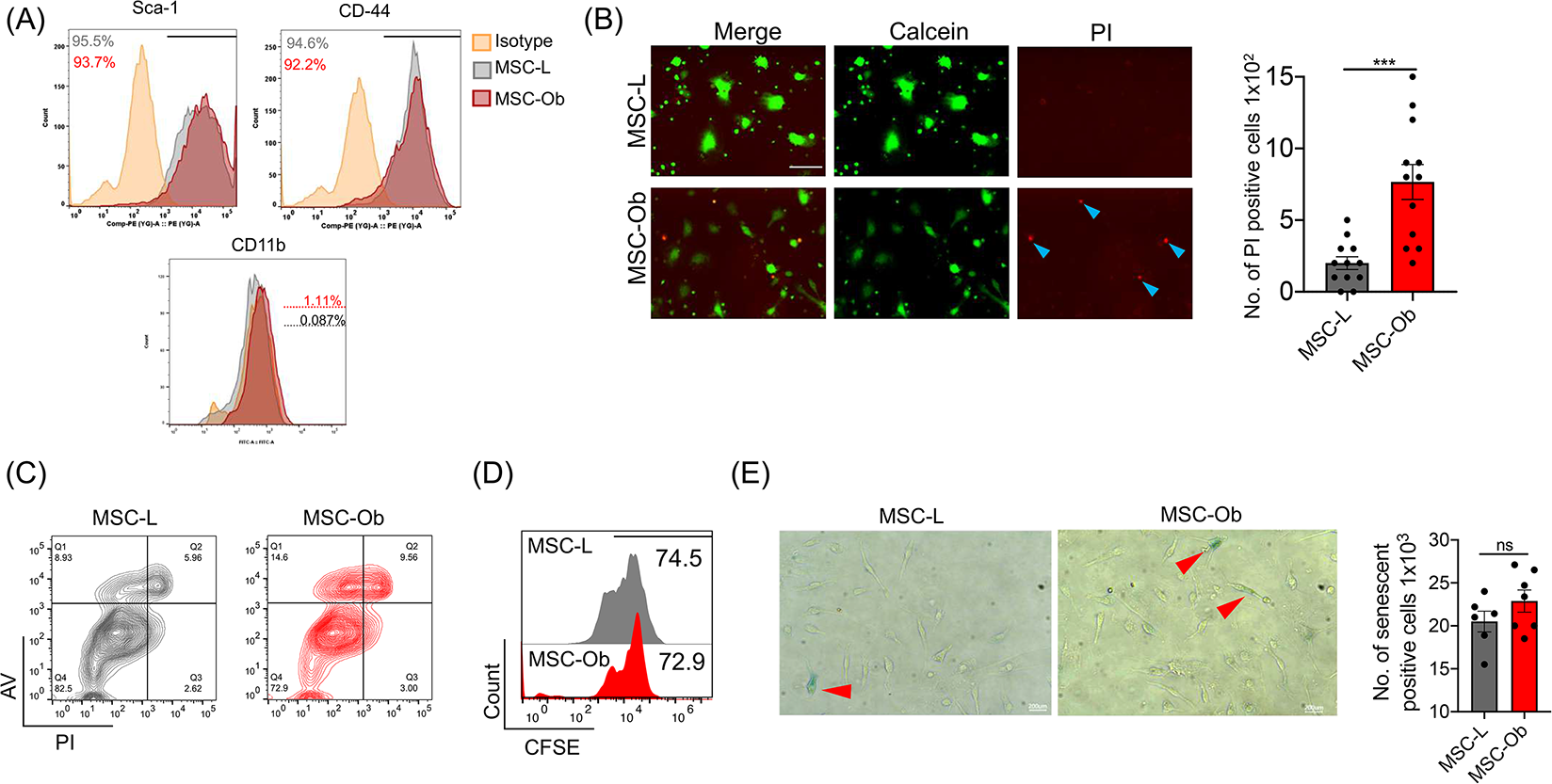
MSC-Ob show cell death under culture conditions with a trend toward an increase in cellular senescence. **(A)** Flow cytometry analysis of stem cell markers expressed by MSC-L and MSC-Ob. (**B**) Representative images show cells stained with propidium iodide (PI) and calcein green. The number of PI-positive cells counted per 100 cells in the images *(B)*. The PI-positive cells indicate cell death (*Right panel*). (**C**) Scatter plot showing the percentage of death measured by staining the cells with PI and annexin V (AV). (**D**) Cell proliferation was measured by staining the cells with CFSE. (**E**) Representative phase-contrast images of cells stained with Senescence-associated β-galactosidase staining (SA-β-gal) are shown in blue. Red arrow heads show the β-gal positive cells. The number of SA-β-gal positive cells were counted and represented as percent senescent cells per 1000 cells counted. Data is shown as Mean±SEM. \*\*\**P* < 0.005; ns (non-significant). Scale bars: B: 50 µm; E: 200 µm.

**Figure S3.**
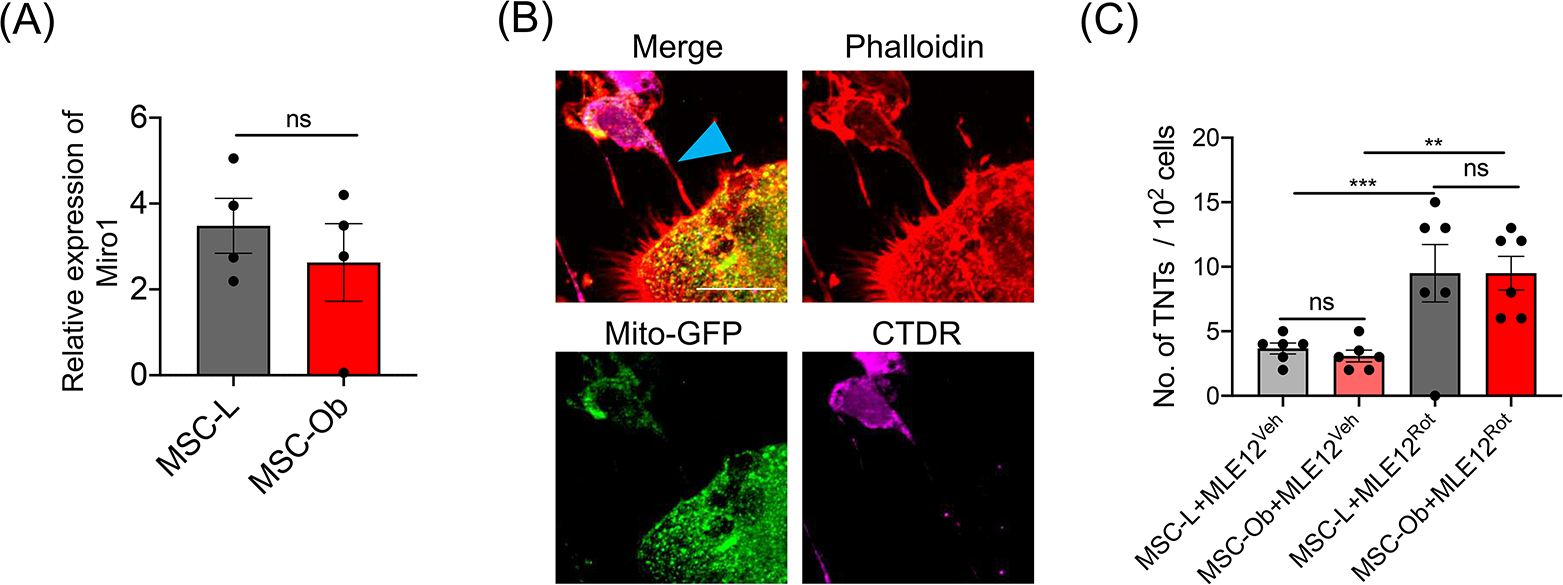
Miro1 expression and TNT formation do not change between MSC-L and MSC-Ob: **(A)** Miro1 RT-qPCR was performed in RNA extracted from the cells and plotted as a relative expression by normalizing with β-actin expression. **(B)** Representative images show tunneling nanotube formation between MSCs and MLE12. MSCs were transduced with mito-GFP (green) and co-cultured with CTDR stained MLE12 for 24 hrs before fixing and staining with Phalloidin (red). The Blue arrowhead shows the TNT formation between MSC and MLE12**. (C)** TNT quantitation between MSCs and MLE12. Data is shown as Mean±SEM. \*\*\*\**P* < 0.001; \*\**P* < 0.001; ns (non-significant). Scale bars: 10 µm.

**Figure S4.**
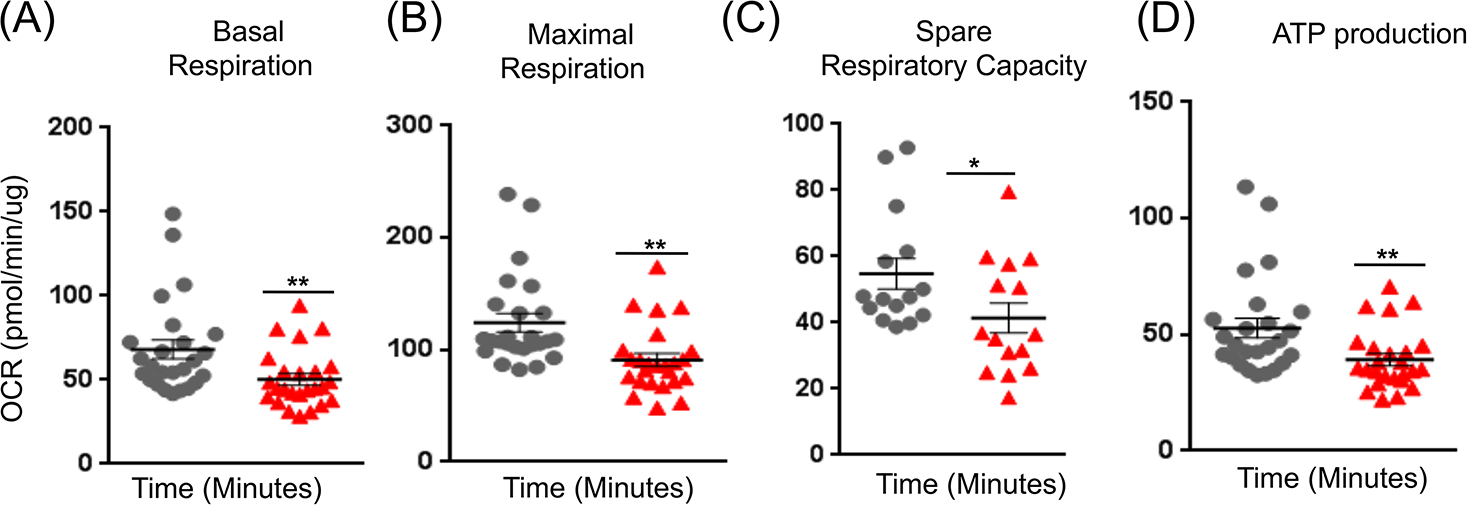
MSC-Ob show decline in mitochondrial bioenergetics: **(A)** Basal respiration was measured by monitoring the oxygen consumption rate (OCR) in MSC-L and MSC-Ob, respectively. (**B-D**) Similarly, maximal respiration, spare respiratory capacity, ATP production was measured. Data is shown as Mean±SEM. \*\**P* < 0.001. \**P* < 0.05.

**Figure S5.**
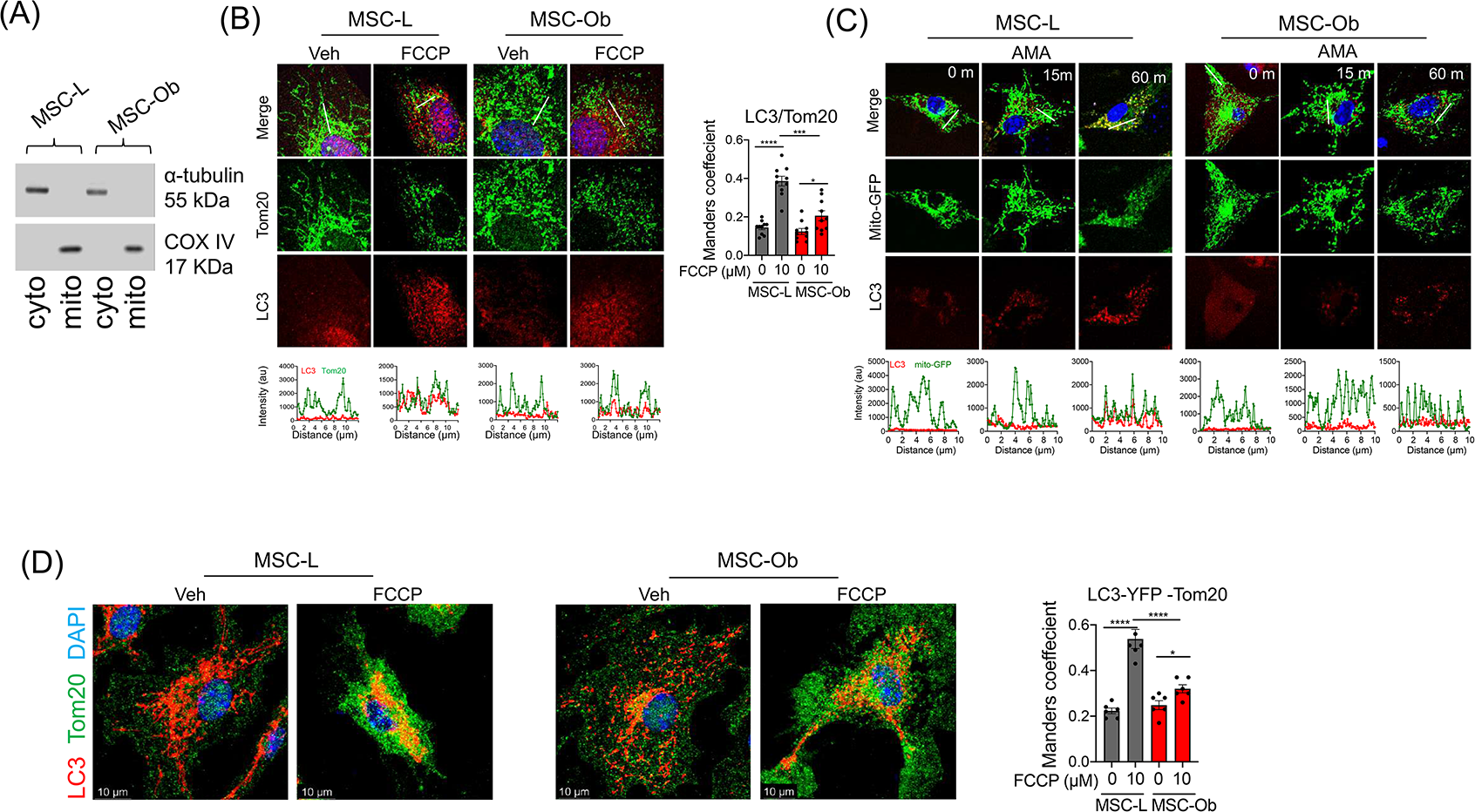
LC3-dependent autophagosome formation is reduced in MSC-Ob. (**A**) Immunoblot of mitochondrial and cytosolic extracts showing α-tubulin expression in cytosolic extract and COX-IV expression in the mitochondrial extract (**B**) Representative images of the cells stained for LC3 (red), Tom20 (green), and DAPI (blue) treated with DMSO (Veh) or FCCP 10µM) for 1 hr. (*Below panel*) Line scan shows colocalization in the regions shown highlighted in images. (*right panel*) show degree of colocalization between LC3 and Tom20 represented as Mander’s coefficient. (**C**) Similarly, cells were treated with 10µM antimycin A (AMA) for 0, 15, and 120 m. The part of this image is also shown in main *Figure 2F*. (**D**) Representative images of cells transduced with LC3-YFP vector (green) and stained for Tom20 (red) and DAPI (blue). The cells were treated with DMSO (Veh) or FCCP (10µM). Mander’s coefficient was calculated to determine the degree of colocalization between LC3-YFP and Tom20 (*right panel*). Data is shown as Mean±SEM. \*\*\*\**P* < 0.001; \*\*\**P* < 0.005; \**P* < 0.05. Scale bars: 10 µm.

**Figure S6.**
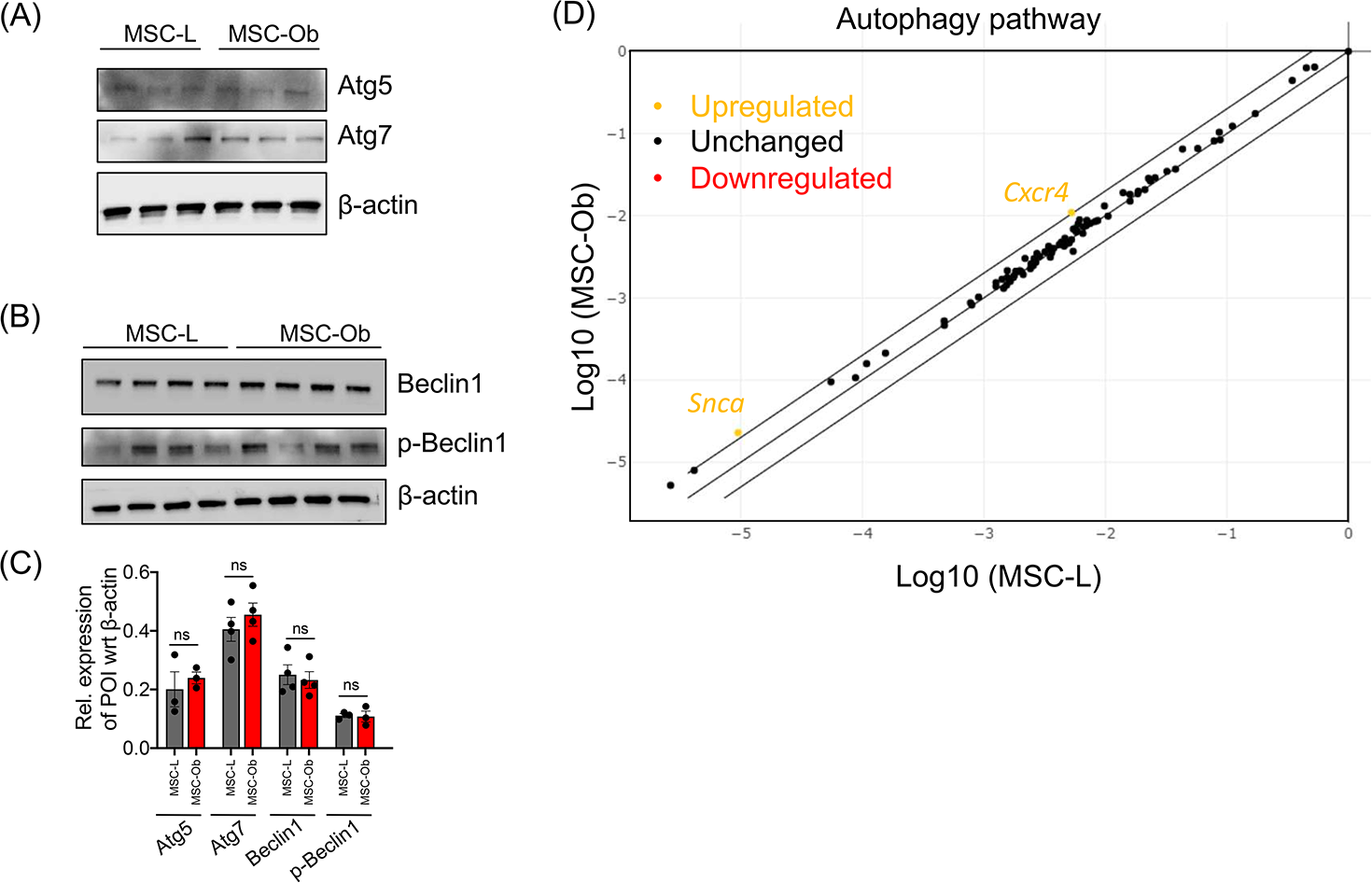
The expression of core autophagy markers is not altered in the MSC-Ob: **(A)** Immunoblots of Atg5, Atg7, and β-actin in the cell lysates prepared from MSCs. (**B**) Similarly, immunoblotting was done for total and phosphorylated forms of Beclin along with β-actin loading as control. (**C**) Densitometry analysis of the blots shown in *A, B*. (**D**) Autophagy pathway analysis was done using the RT2 profiler PCR array. The upregulated genes are shown in yellow dots, and black dots represent genes that did not significantly change expression. Data is shown as Mean±SEM. ns (non-significant).

**Figure S7.**
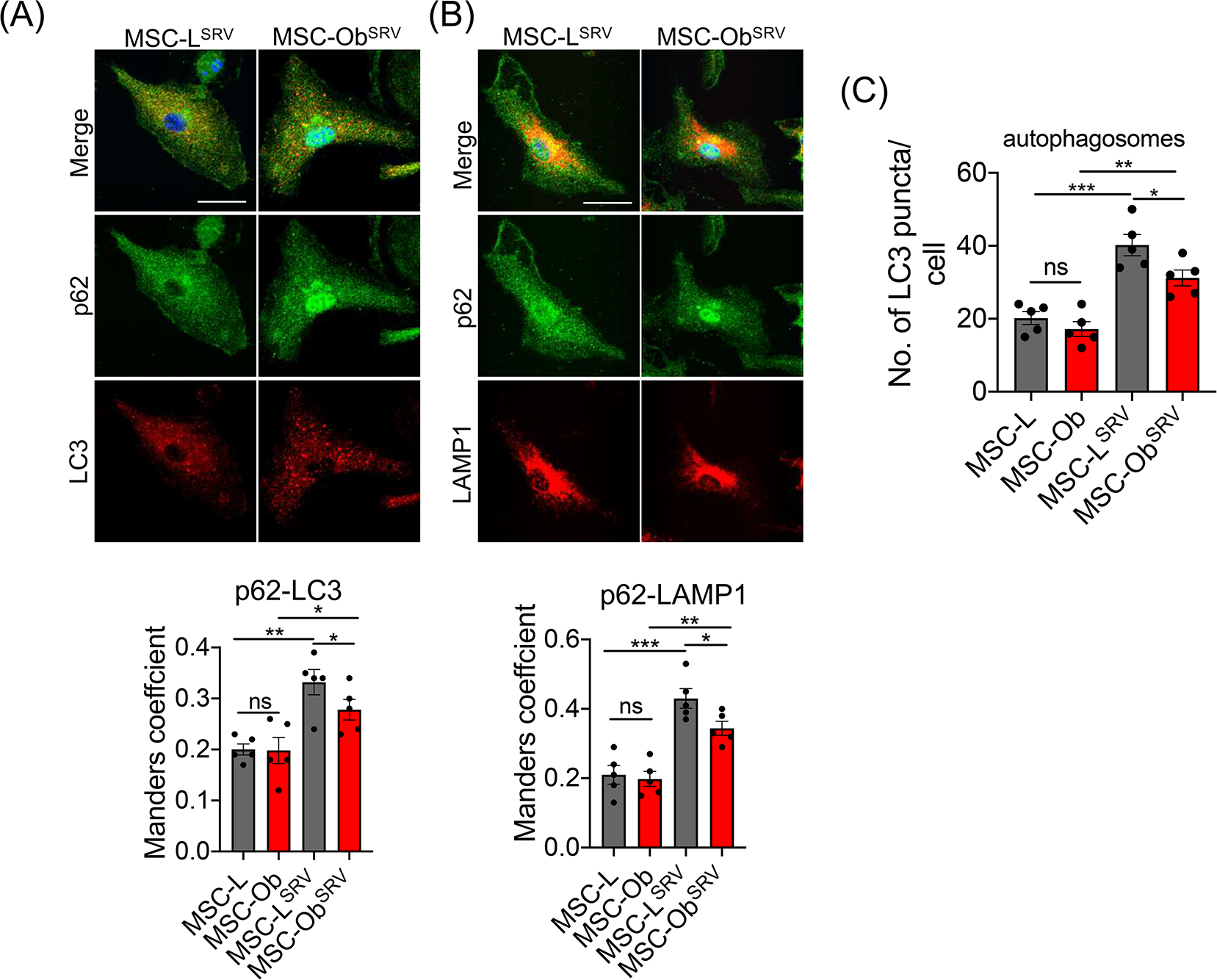
Starvation induces autophagosome formation and autophagy of p62: **(A)** Representative images of p62 colocalization with LC3 in cells cultured under normal conditions and starvation (SRV). (**B**) Similarly, cells were stained with lysotracker deep red (LTDR). The degree of colocalization was calculated between P62 with LC3 or LTDR, respectively (below panels). (**C**) The autophagosome formation was calculated in cells stained for LC3 under normal and SRV conditions. Data is shown as Mean±SEM. \*\*\**P* < 0.005; \*\**P* < 0.01; \**P* < 0.05; ns (non-significant). Scale bars: 10 µm.

**Figure S8.**
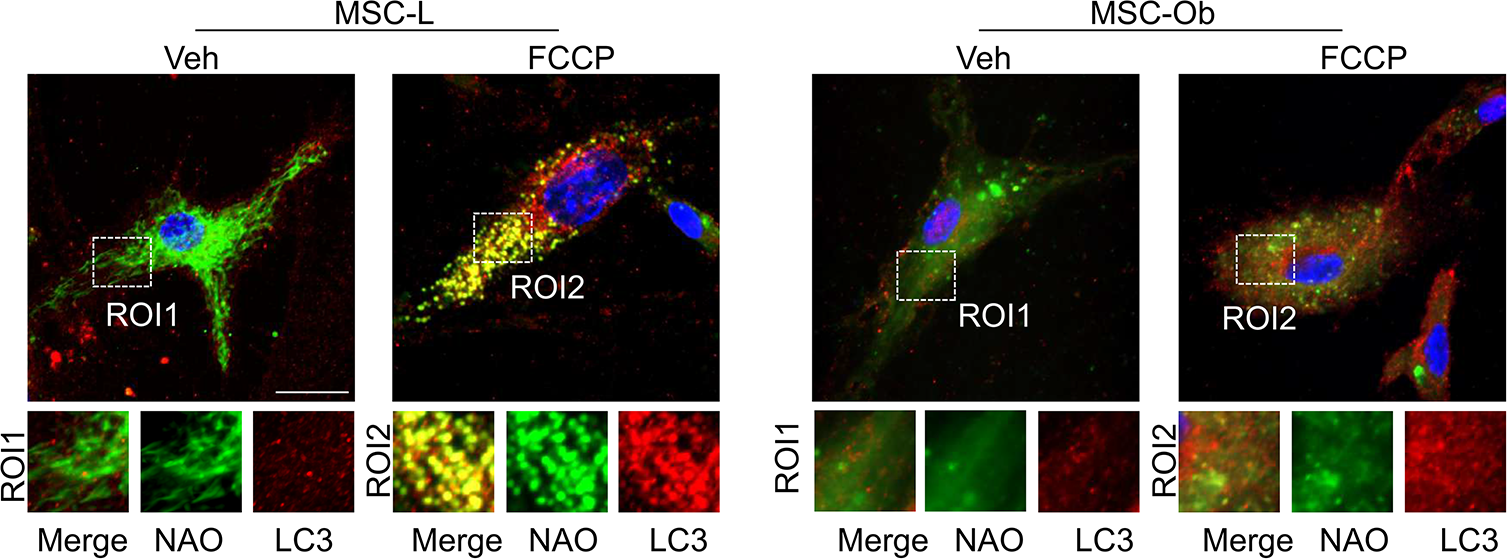
Reduced cardiolipin colocalization with LC3: The representative images correspond to the insets shown in *Figure 3L.* To induce depolarization, MSC-L and MSC-Ob were treated with Veh or FCCP and stained for NAO (green) and LC3 (red). The insets below show the region of interest (ROI). Scale bars: 10 µm.

**Figure S9.**
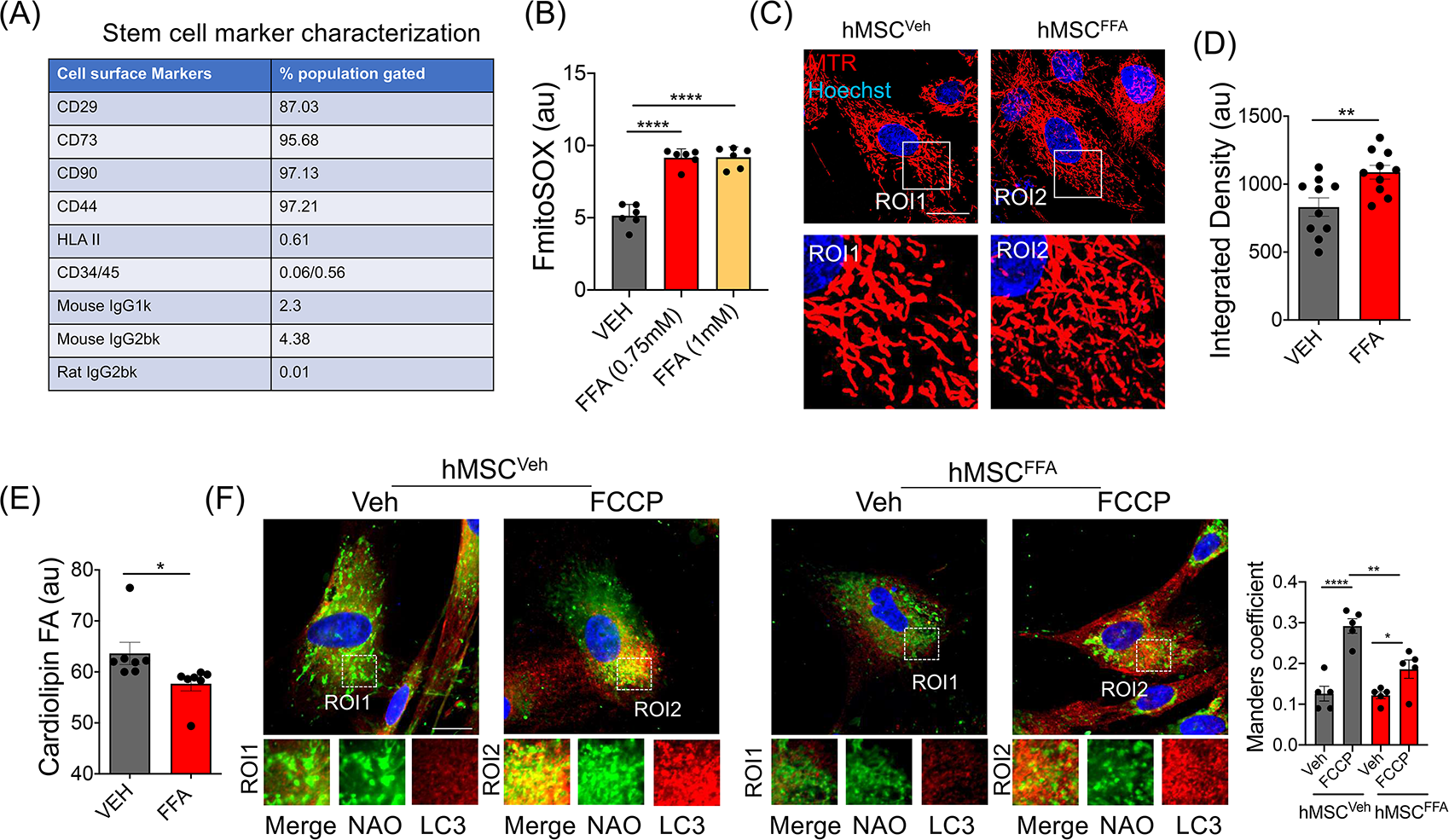
Human MSCs show mitochondrial dysfunction and reduced cardiolipin content upon FFA treatment: **(A)** hMSCs isolated from the bone marrow of normal healthy individuals were analyzed for the expression of stem cell markers. Flow cytometry was performed, and the data is represented as % expression with respect to the isotype control. **(B)** mtROS was measured in hMSC treated with Veh (DMSO) or free fatty acids (FFA) for 24 hrs. The analysis was done by flow cytometry using mitoSOX red. (**C**) Representative images of hMSCs stained with MTR (red) and Hoechst (blue) to find the changes in mitochondrial morphology. (**D**) Mitochondrial mass was calculated in the cells (*C*) stained with MTR. (**E**) Cardiolipin content was measured in hMSCs by flow cytometry and represented as the bar graph. (**F**) Representative images of cells stained with NAO (green) and LC3 (red). The cells were treated with Veh or FCCP (µM) for 1 hr before imaging. (*Right panel*) shows corresponding Mander’s coefficient of the images shown in *F*. Data is shown as Mean±SEM. \*\*\*\**P* < 0.001; \*\**P* < 0.01; \**P* < 0.05. Scale bars: 10 µm.

**Figure S10.**
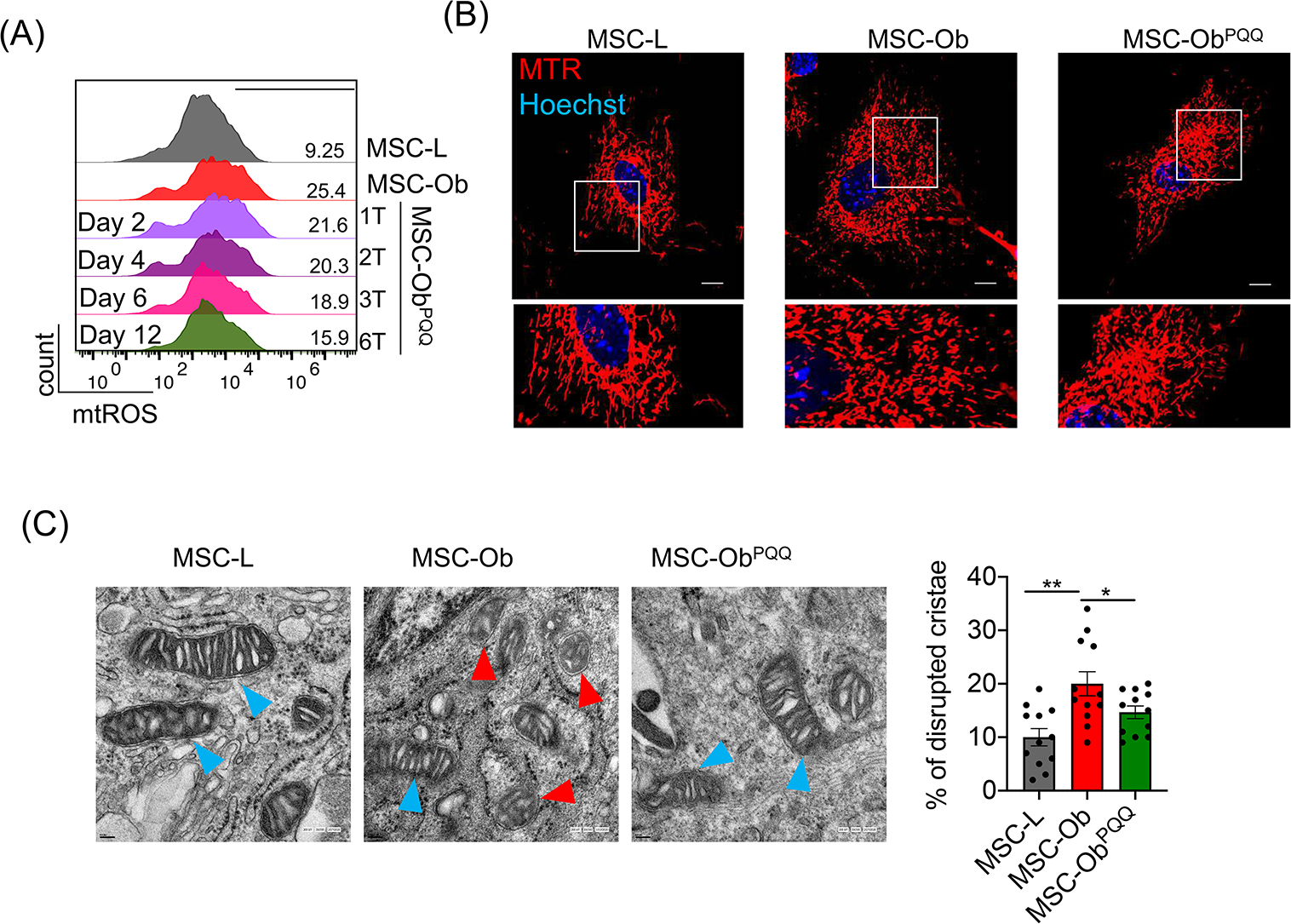
PQQ treatments restore mitochondrial function and ultrastructural changes: (**A**) Representative histograms of cells showing mtROS. MSC-Ob were treated with PQQ (30µM) for the indicated days with media change after every 2 days and supplemented with fresh PQQ (shown as T: no. of treatments). (**B**) Representative images show changes in mitochondrial shape and size. (**C**) Representative EM images showing mitochondrial ultrastructural changes with disrupted cristae indicated by red arrow head and healthy mitochondria by blue arrow head. The number of cristae were quantified and presented as percentage disrupted cristae per 100 cristae counted (*right panel*). Data is shown as Mean±SEM. \*\**P* < 0.01; \**P* < 0.05. Scale bars: C: 10 µm; D: 0.1µm.

**Figure S11.**
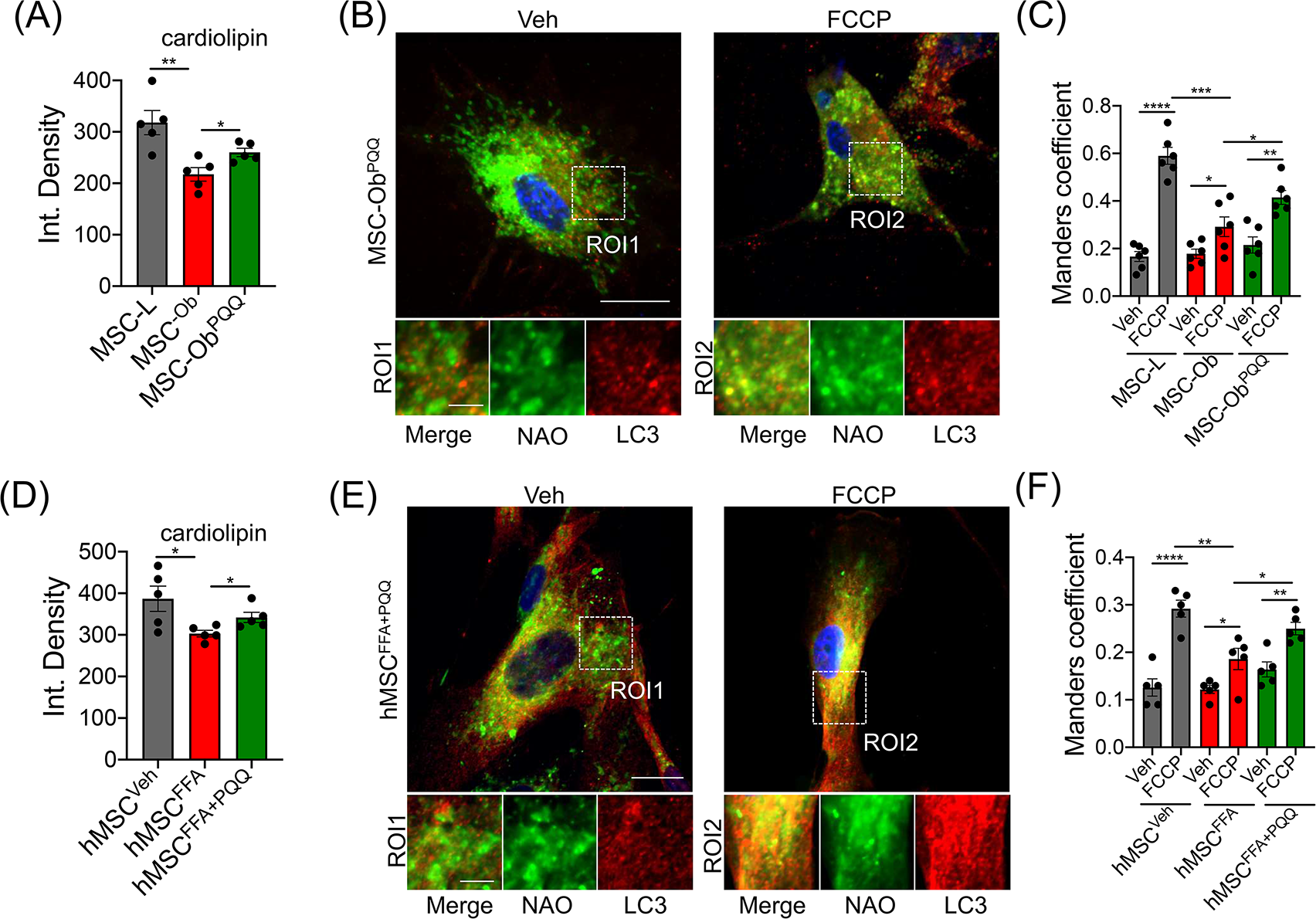
PQQ restores cardiolipin and LC3 colocalization: (**A**) Bar graph showing the signal quantitation of NAO (represents cardiolipin content) in MSCs treated with Veh or PQQ. (**B**) Representative images showing colocalization of cardiolipin stained with NAO (green) with LC3 (red) in cells treated with Veh or FCCP (10 µM for 1 hr.). The insets below show the respective ROIs of the square dotted area. (**C**) The corresponding Mander’s coefficient of the images shows the extent of colocalization. The histogram panel *C* is same as shown in the *Figure 3L* which is without PQQ data. (**D**) Bar graph showing the signal quantitation of NAO (represents cardiolipin content) in hMSCs treated with Veh, FFA and FFA+PQQ (**E**) Representative images of hMSCs treated with PQQ and stained with NAO (green) and LC3 (red) with insets below the respective ROIs. (**F**) Corresponding Mander’s coefficient representation, the extent of colocalization between cardiolipin and LC3. Data is shown as Mean±SEM. \*\*\*\**P* < 0.001; \*\*\**P* < 0.005; \*\**P* < 0.01; \**P* < 0.05. Scale bars: C: 10 µm; Insets: 5µm.

**Figure S12.**
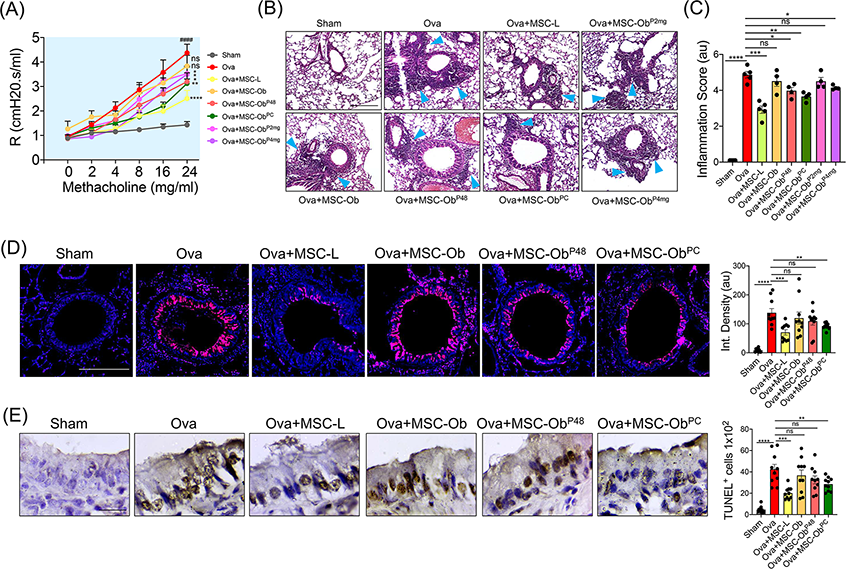
MSC-Ob treated with PQQ restore airway mechanics, airway remodeling, and airway epithelial cell damage: (**A**) AHR in Ova-induced allergic airway inflammation model transplanted with MSCs obtained from lean or HFD mice and treated with various concentrations and time points of PQQ. The groups 2mg and 4mg refer to the HFD mice which were fed PQQ for 15 days before harvesting MSCs. (**B**) H&E images showing the airway infiltration of cells (blue arrowheads). (**C**) Inflammatory score representing the extent of airway inflammatory cell infiltration. (**D**) Representative images of tissue sections stained with PAS and pseudocolored. The pink color represents the mucus secretion, while the blue color is the representation of nuclei. The images were subjected to quantitative analysis and represented as integrated density *(right panel)*. (**E**) Representative images showing TUNEL staining in bronchial epithelial cells. Brown represents the TUNEL positive cells (cell death) per 100 nuclei counted stained with hematoxylin (blue). Mean±SEM. \*\*\*\**P* < 0.001; \*\*\**P* < 0.005; \*\**P* < 0.01; \**P* < 0.05; ns (non-significant). Scale bars: B: 200 µm; D: 100 µm; E: 50 µm.

## Reference

1. M. F. Pittenger, et al., Mesenchymal stem cell perspective: cell biology to clinical progress. NPJ Regen Med. 4, 22 (2019).

2. O. Levy et al., Shattering barriers toward clinically meaningful MSC therapies. Sci Adv. 6, eaba6884 (2020).

3. A. N. Lau, M. Goodwin, C. F. Kim, D. J. Weiss, Stem cells and regenerative medicine in lung biology and diseases. Mol Ther. 20, 1116–1130 (2012).

4. A. K. Shetty, P. A. Shetty, G. Zanirati, K. Jin, Further validation of the efficacy of mesenchymal stem cell infusions for reducing mortality in COVID-19 patients with ARDS. NPJ Regen Med. 6, 53 (2021).

5. K. Nemeth et al., Bone marrow stromal cells use TGF-beta to suppress allergic responses in a mouse model of ragweed-induced asthma. Proc Natl Acad Sci U S A. 107, 5652–5657 (2010).

6. S. Rani, A. E. Ryan, M. D. Griffin, T. Ritter, Mesenchymal Stem Cell-derived Extracellular Vesicles: Toward Cell-free Therapeutic Applications. Mol Ther. 23, 812–823 (2015).

7. A. R. R. Weiss, M. H. Dahlke, Immunomodulation by Mesenchymal Stem Cells (MSCs): Mechanisms of Action of Living, Apoptotic, and Dead MSCs. Front Immunol. 10, 1191 (2019).

8. T. Ahmad et al., Miro1 regulates intercellular mitochondrial transport & enhances mesenchymal stem cell rescue efficacy. Embo j. 33, 994–1010 (2014).

9. M. O. Gomzikova, V. James, A. A. Rizvanov, Mitochondria Donation by Mesenchymal Stem Cells: Current Understanding and Mitochondria Transplantation Strategies. Front Cell Dev Biol. 9, 653322 (2021).

10. D. Liu et al., Intercellular mitochondrial transfer as a means of tissue revitalization. Signal Transduct Target Ther. 6, 65 (2021).

11. D. Jiang et al., Mitochondrial transfer of mesenchymal stem cells effectively protects corneal epithelial cells from mitochondrial damage. Cell Death Dis. 7, e2467 (2016).

12. J. Gao et al., Endoplasmic reticulum mediates mitochondrial transfer within the osteocyte dendritic network. Sci Adv. 5, eaaw7215 (2019).

13. M. N. Islam et al., Mitochondrial transfer from bone-marrow-derived stromal cells to pulmonary alveoli protects against acute lung injury. Nat Med. 18, 759–765 (2012).

14. T. Huang et al., Iron oxide nanoparticles augment the intercellular mitochondrial transfer-mediated therapy. Sci Adv. 7, eabj0534 (2021).

15. K. Saito et al., Exogenous mitochondrial transfer and endogenous mitochondrial fission facilitate AML resistance to OxPhos inhibition. Blood Adv. 5, 4233–4255 (2021).

16. R. Moschoi et al., Protective mitochondrial transfer from bone marrow stromal cells to acute myeloid leukemic cells during chemotherapy. Blood. 128, 253–264 (2016).

17. R. Burt et al., Activated stromal cells transfer mitochondria to rescue acute lymphoblastic leukemia cells from oxidative stress. Blood. 134, 1415–1429 (2019).

18. Y. Yao et al., Connexin 43-Mediated Mitochondrial Transfer of iPSC-MSCs Alleviates Asthma Inflammation. Stem Cell Reports. 11, 1120–1135 (2018).

19. X. Li et al., Mitochondrial transfer of induced pluripotent stem cell-derived mesenchymal stem cells to airway epithelial cells attenuates cigarette smoke-induced damage. Am J Respir Cell Mol Biol. 51, 455–465 (2014).

20. A. C. Court et al., Mitochondrial transfer from MSCs to T cells induces Treg differentiation and restricts inflammatory response. EMBO Rep. 21, e48052 (2020).

21. Y. Meng et al., Obesity-induced mitochondrial dysfunction in porcine adipose tissue-derived mesenchymal stem cells. J Cell Physiol. 233, 5926–5936 (2018).

22. K. Kornicka, J. Houston, K. Marycz, Dysfunction of Mesenchymal Stem Cells Isolated from Metabolic Syndrome and Type 2 Diabetic Patients as Result of Oxidative Stress and Autophagy may Limit Their Potential Therapeutic Use. Stem Cell Rev Rep. 14, 337–345 (2018).

23. L. M. Pérez et al., Altered metabolic and stemness capacity of adipose tissue-derived stem cells from obese mouse and human. PLoS One. 10, e0123397 (2015).

24. Y. Li et al., GSK3 inhibitor ameliorates steatosis through the modulation of mitochondrial dysfunction in hepatocytes of obese patients. iScience. 24, 102149 (2021).

25. O. Kizilay Mancini et al., Mitochondrial Oxidative Stress Reduces the Immunopotency of Mesenchymal Stromal Cells in Adults With Coronary Artery Disease. Circ Res. 122, 255–266 (2018).

26. M. S. Choudhery, M. Badowski, A. Muise, J. Pierce, D. T. Harris, Donor age negatively impacts adipose tissue-derived mesenchymal stem cell expansion and differentiation. J Transl Med. 12, 8 (2014).

27. K. Kornicka, K. Marycz, K. A. Tomaszewski, M. Marędziak, A. Śmieszek, The Effect of Age on Osteogenic and Adipogenic Differentiation Potential of Human Adipose Derived Stromal Stem Cells (hASCs) and the Impact of Stress Factors in the Course of the Differentiation Process. Oxid Med Cell Longev. 2015, 309169 (2015).

28. X. Ye et al., Age-Related Changes in the Regenerative Potential of Adipose-Derived Stem Cells Isolated from the Prominent Fat Pads in Human Lower Eyelids. PLoS One. 11, e0166590 (2016).

29. A. Stolzing, E. Jones, D. McGonagle, A. Scutt, Age-related changes in human bone marrow-derived mesenchymal stem cells: consequences for cell therapies. Mech Ageing Dev. 129, 163–173 (2008).

30. Y. Zhang et al., Adult mesenchymal stem cell ageing interplays with depressed mitochondrial Ndufs6. Cell Death Dis. 11, 1075 (2020).

31. K. Marycz, K. Kornicka, K. Basinska, A. Czyrek, Equine Metabolic Syndrome Affects Viability, Senescence, and Stress Factors of Equine Adipose-Derived Mesenchymal Stromal Stem Cells: New Insight into EqASCs Isolated from EMS Horses in the Context of Their Aging. Oxid Med Cell Longev. 2016, 4710326 (2016).

32. Y. Li et al., Metabolic syndrome increases senescence-associated micro-RNAs in extracellular vesicles derived from swine and human mesenchymal stem/stromal cells. Cell Commun Signal. 18, 124 (2020).

33. C. Serena et al., Obesity and Type 2 Diabetes Alters the Immune Properties of Human Adipose Derived Stem Cells. Stem Cells. 34, 2559–2573 (2016).

34. R. J. Youle, D. P. Narendra, Mechanisms of mitophagy. Nat Rev Mol Cell Biol. 12, 9–14 (2011).

35. R. J. Youle, Mitochondria-Striking a balance between host and endosymbiont. Science. 365, (2019).

36. S. Pickles, P. Vigié, R. J. Youle, Mitophagy and Quality Control Mechanisms in Mitochondrial Maintenance. Curr Biol. 28, R170–r185 (2018).

37. D. A. Kubli, B. Gustafsson Å, Mitochondria and mitophagy: the yin and yang of cell death control. Circ Res. 111, 1208–1221 (2012).

38. M. A. Lampert et al., BNIP3L/NIX and FUNDC1-mediated mitophagy is required for mitochondrial network remodeling during cardiac progenitor cell differentiation. Autophagy. 15, 1182–1198 (2019).

39. S. von Stockum, E. Marchesan, E. Ziviani, Mitochondrial quality control beyond PINK1/Parkin. Oncotarget. 9, 12550–12551 (2018).

40. P. Terešak et al., Regulation of PRKN-independent mitophagy. Autophagy. 1–16 (2021).

41. F. Zhang et al., P53 and Parkin co-regulate mitophagy in bone marrow mesenchymal stem cells to promote the repair of early steroid-induced osteonecrosis of the femoral head. Cell Death Dis. 11, 42 (2020).

42. D. G. Phinney et al., Mesenchymal stem cells use extracellular vesicles to outsource mitophagy and shuttle microRNAs. Nat Commun. 6, 8472 (2015).

43. J. H. Lee, Y. M. Yoon, K. H. Song, H. Noh, S. H. Lee, Melatonin suppresses senescence-derived mitochondrial dysfunction in mesenchymal stem cells via the HSPA1L-mitophagy pathway. Aging Cell. 19, e13111 (2020).

44. L. M. Pérez, A. Bernal, N. San Martín, B. G. Gálvez, Obese-derived ASCs show impaired migration and angiogenesis properties. Arch Physiol Biochem. 119, 195–201 (2013).

45. C. L. Wu, B. O. Diekman, D. Jain, F. Guilak, Diet-induced obesity alters the differentiation potential of stem cells isolated from bone marrow, adipose tissue and infrapatellar fat pad: the effects of free fatty acids. Int J Obes (Lond). 37, 1079–1087 (2013).

46. S. Ceccariglia, A. Cargnoni, A. R. Silini, O. Parolini, Autophagy: a potential key contributor to the therapeutic action of mesenchymal stem cells. Autophagy. 16, 28–37 (2020).

47. V. P. Singh et al., Metabolic Syndrome Is Associated with Increased Oxo-Nitrative Stress and Asthma-Like Changes in Lungs. PLoS One. 10, e0129850 (2015).

48. F. Wang et al., MICAL2PV suppresses the formation of tunneling nanotubes and modulates mitochondrial trafficking. EMBO Rep. 22, e52006 (2021).

49. A. Luciani et al., Impaired mitophagy links mitochondrial disease to epithelial stress in methylmalonyl-CoA mutase deficiency. Nat Commun. 11, 970 (2020).

50. D. Narendra, J. E. Walker, R. Youle, Mitochondrial quality control mediated by PINK1 and Parkin: links to parkinsonism. Cold Spring Harb Perspect Biol. 4, (2012).

51. T. Ahmad et al., Computational classification of mitochondrial shapes reflects stress and redox state. Cell Death Dis. 4, e461 (2013).

52. B. P. Festa et al., Impaired autophagy bridges lysosomal storage disease and epithelial dysfunction in the kidney. Nat Commun. 9, 161 (2018).

53. B. C. Hammerling et al., A Rab5 endosomal pathway mediates Parkin-dependent mitochondrial clearance. Nat Commun. 8, 14050 (2017).

54. Y. Nishida et al., Discovery of Atg5/Atg7-independent alternative macroautophagy. Nature. 461, 654–658 (2009).

55. Y. Ichimura, E. Kominami, K. Tanaka, M. Komatsu, Selective turnover of p62/A170/SQSTM1 by autophagy. Autophagy. 4, 1063–1066 (2008).

56. S. C. da Silva Rosa et al., BNIP3L/Nix-induced mitochondrial fission, mitophagy, and impaired myocyte glucose uptake are abrogated by PRKA/PKA phosphorylation. Autophagy. 17, 2257–2272 (2021).

57. C. T. Chu, H. Bayır, V. E. Kagan, LC3 binds externalized cardiolipin on injured mitochondria to signal mitophagy in neurons: implications for Parkinson disease. Autophagy. 10, 376–378 (2014).

58. C. T. Chu et al., Cardiolipin externalization to the outer mitochondrial membrane acts as an elimination signal for mitophagy in neuronal cells. Nat Cell Biol. 15, 1197–1205 (2013).

59. H. Malhi, S. F. Bronk, N. W. Werneburg, G. J. Gores, Free fatty acids induce JNK-dependent hepatocyte lipoapoptosis. J Biol Chem. 281, 12093–12101 (2006).

60. W. Chowanadisai et al., Pyrroloquinoline quinone stimulates mitochondrial biogenesis through cAMP response element-binding protein phosphorylation and increased PGC-1alpha expression. J Biol Chem. 285, 142–152 (2010).

61. Q. Zhang et al., Pyrroloquinoline Quinone Inhibits Rotenone-Induced Microglia Inflammation by Enhancing Autophagy. Molecules. 25, (2020).

62. Q. Cheng et al., Pyrroloquinoline quinone promotes mitochondrial biogenesis in rotenone-induced Parkinson’s disease model via AMPK activation. Acta Pharmacol Sin. 42, 665–678 (2021).

63. X. Li et al., Mesenchymal stem cells alleviate oxidative stress-induced mitochondrial dysfunction in the airways. J Allergy Clin Immunol. 141, 1634–1645.e1635 (2018).

64. Z. Z. Liu et al., Autophagy receptor OPTN (optineurin) regulates mesenchymal stem cell fate and bone-fat balance during aging by clearing FABP3. Autophagy. 17, 2766–2782 (2021).

65. J. B. Spinelli, M. C. Haigis, The multifaceted contributions of mitochondria to cellular metabolism. Nat Cell Biol. 20, 745–754 (2018).

66. S. Paliwal, R. Chaudhuri, A. Agrawal, S. Mohanty, Human tissue-specific MSCs demonstrate differential mitochondria transfer abilities that may determine their regenerative abilities. Stem Cell Res Ther. 9, 298 (2018).

67. Y. Kuang et al., Structural basis for the phosphorylation of FUNDC1 LIR as a molecular switch of mitophagy. Autophagy. 12, 2363–2373 (2016).

68. M. Marinković, M. Šprung, I. Novak, Dimerization of mitophagy receptor BNIP3L/NIX is essential for recruitment of autophagic machinery. Autophagy. 17, 1232–1243 (2021).

69. M. Onishi, K. Yamano, M. Sato, N. Matsuda, K. Okamoto, Molecular mechanisms and physiological functions of mitophagy. Embo j. 40, e104705 (2021).

70. H. Wu et al., Deficiency of mitophagy receptor FUNDC1 impairs mitochondrial quality and aggravates dietary-induced obesity and metabolic syndrome. Autophagy. 15, 1882–1898 (2019).

71. C. G. Towers et al., Mitochondrial-derived vesicles compensate for loss of LC3-mediated mitophagy. Dev Cell. 56, 2029–2042.e2025 (2021).

72. R. Tao et al., Pyrroloquinoline quinone preserves mitochondrial function and prevents oxidative injury in adult rat cardiac myocytes. Biochem Biophys Res Commun. 363, 257–262 (2007).

73. X. Yuan, T. M. Logan, T. Ma, Metabolism in Human Mesenchymal Stromal Cells: A Missing Link Between hMSC Biomanufacturing and Therapy? Front Immunol. 10, 977 (2019).

74. N. Konari, K. Nagaishi, S. Kikuchi, M. Fujimiya, Mitochondria transfer from mesenchymal stem cells structurally and functionally repairs renal proximal tubular epithelial cells in diabetic nephropathy in vivo. Sci Rep. 9, 5184 (2019).

75. J. Levoux et al., Platelets Facilitate the Wound-Healing Capability of Mesenchymal Stem Cells by Mitochondrial Transfer and Metabolic Reprogramming. Cell Metab. 33, 283–299.e289 (2021).

76. J. R. Brestoff et al., Intercellular Mitochondria Transfer to Macrophages Regulates White Adipose Tissue Homeostasis and Is Impaired in Obesity. Cell Metab. 33, 270–282.e278 (2021).

77. X. L. Fan, Y. Zhang, X. Li, Q. L. Fu, Mechanisms underlying the protective effects of mesenchymal stem cell-based therapy. Cell Mol Life Sci. 77, 2771–2794 (2020).

78. R. A. Denu, P. Hematti, Effects of Oxidative Stress on Mesenchymal Stem Cell Biology. Oxid Med Cell Longev. 2016, 2989076 (2016).

79. G. Ye et al., Oxidative stress-mediated mitochondrial dysfunction facilitates mesenchymal stem cell senescence in ankylosing spondylitis. Cell Death Dis. 11, 775 (2020).

80. A. J. C. Bloor et al., Production, safety and efficacy of iPSC-derived mesenchymal stromal cells in acute steroid-resistant graft versus host disease: a phase I, multicenter, open-label, dose-escalation study. Nat Med. 26, 1720–1725 (2020).

81. K. Sun, X. Jing, J. Guo, X. Yao, F. Guo, Mitophagy in degenerative joint diseases. Autophagy. 17, 2082–2092 (2021).

82. H. Zhu et al., A protocol for isolation and culture of mesenchymal stem cells from mouse compact bone. Nat Protoc. 5, 550–560 (2010).

83. S. B. Nandy, S. Mohanty, M. Singh, M. Behari, B. Airan, Fibroblast Growth Factor-2 alone as an efficient inducer for differentiation of human bone marrow mesenchymal stem cells into dopaminergic neurons. J Biomed Sci. 21, 83 (2014).

84. R. K. Motiani et al., STIM1 activation of adenylyl cyclase 6 connects Ca(2+) and cAMP signaling during melanogenesis. Embo j. 37, (2018).

85. W. Stauffer, H. Sheng, H. N. Lim, EzColocalization: An ImageJ plugin for visualizing and measuring colocalization in cells and organisms. Sci Rep. 8, 15764 (2018).

86. T. Ahmad et al., Impaired mitophagy leads to cigarette smoke stress-induced cellular senescence: implications for chronic obstructive pulmonary disease. Faseb j. 29, 2912–2929 (2015).

87. J. P. Rooney et al., PCR based determination of mitochondrial DNA copy number in multiple species. Methods Mol Biol. 1241, 23–38 (2015).

88. T. Ahmad et al., Simvastatin improves epithelial dysfunction and airway hyperresponsiveness: from asymmetric dimethyl-arginine to asthma. Am J Respir Cell Mol Biol. 44, 531–539 (2011).

## Reference

1. D. Ryu et al., Urolithin A induces mitophagy and prolongs lifespan in C. elegans and increases muscle function in rodents. Nat. Med. 22, 879–888 (2016).

2. Y. Choi et al., Enhancement of Mesenchymal Stem Cell-Driven Bone Regeneration by Resveratrol-Mediated SOX2 Regulation. Aging Dis. 10, 818–833 (2019).

3. Y. Zhang et al., Adult mesenchymal stem cell ageing interplays with depressed mitochondrial Ndufs6. Cell Death Dis. 11, 1075 (2020).

4. W. Chowanadisai et al., Pyrroloquinoline quinone stimulates mitochondrial biogenesis through cAMP response element-binding protein phosphorylation and increased PGC-1alpha expression. J. Biol. Chem. 285, 142–152 (2010).

5. E. F. Fang et al., NAD(+) augmentation restores mitophagy and limits accelerated aging in Werner syndrome. Nat Commun 10, 5284 (2019).

6. Y. T. Wu et al., Dual role of 3-methyladenine in modulation of autophagy via different temporal patterns of inhibition on class I and III phosphoinositide 3-kinase. J. Biol. Chem. 285, 10850–10861 (2010).

7. C. J. Li, L. Y. Sun, C. Y. Pang, Synergistic protection of N-acetylcysteine and ascorbic acid 2-phosphate on human mesenchymal stem cells against mitoptosis, necroptosis and apoptosis. Sci. Rep. 5, 9819 (2015).

